# Initial Sequencing and Characterization of Gastrointestinal and Oral Microbiota in Urban Pakistani Adults Reveals Abnormally High Levels of Potentially Starch Metabolizing Bacteria in the General Population

**DOI:** 10.1101/419598

**Authors:** Maria Batool, Syed Baqir Ali, Ali Jaan, Kehkishan Khalid, Syeda Aba Ali, Kainat Kamal, Afraz Ahmed Raja, Farzana Gul, Inti Pedroso, Zachary Apte, Arshan Nasir

**Affiliations:** Department of Biosciences, COMSATS University Islamabad, Park Road, Tarlai Kalan, Islamabad 45550, Pakistan; uBiome, San Francisco, CA, United States

## Abstract

We describe the characterization of the gastrointestinal tract (gut) and oral microbiota (bacteria) in 32 urban Pakistani adults representing seven major geographies and six ethnicities in the country. Study participants were between ages 18 and 40, had body mass index between 18 and 25 Kg/m^2^, and were early-career students or professionals belonging to 25 major cities of the country. These individuals donated a total of 61 samples (32 gut and 29 oral) that were subjected to 16S ribosomal RNA (rRNA) gene sequencing. Microbiome composition of Pakistani individuals was compared against the uBiome database of selected individuals who self-reported to be in excellent health. Using the crude measure of percentage overlap or similarity between the gut microbiota profile of Pakistani and uBiome dataset as proxy for health, our sequencing indicated that the Pakistani gut microbiota was moderately healthy relative to the uBiome dataset and Pakistani women appeared healthier relative to men. The Pakistani gut microbiome seemed susceptible to obesity and weight gain, levels of probiotics was very high likely due to the popularity of milk-based and fermented foods in the Pakistani diet, and bacteria that metabolize starch and carbohydrates (typically seen in the gut microbiota of honey bee) were abnormally enriched in the gut of Pakistani men. Our investigations reveal serious issues with the dietary habits and lifestyle of Pakistani individuals of consuming food enriched in high carbohydrates and fats, overcooked in oil and spices, following a sedentary lifestyle, little or no daily intake of fresh fruits, over-consumption of antibiotics from a very early age, and health and hygiene standards that do not meet international standards. Our sequencing is the first step towards generating a country-wide understanding of the impact of the local diet and lifestyle on Pakistani gut microbiota and can help understand its overall association with health and wellness.

## INTRODUCTION

The human body hosts millions of microorganisms including Bacteria, Archaea, Viruses, Fungi, and unicellular Protists that reside on various human body sites such as the gastrointestinal tract (gut), oral cavity (mouth), skin, and others [1]. According to latest estimates, the number of microbial cells in and on the human body is roughly equal to the number of total human cells (∼40 trillion) [2]. The gastrointestinal tract harbors the largest and most diverse ecosystem of human-associated microorganisms in the human body [3]. The human-associated microorganisms, or collectively the human microbiota, forms a strong symbiotic relationship with the human host [4] and is actively involved in a variety of biological processes such as fermentation and digestion of undigested carbohydrates [5], immune system maturation [6–8], and (even) influencing our social behavior [9] through the “gut-brain” axis [10,11]. In turn, abnormal microbiota composition or dysbiosis has been associated with various diseases ranging from metabolic disorders [12,13] to social anxiety and depression [10,14] highlighting the need to maintain a healthy microbial community for human wellness [15]. Indeed, microbiota transplant from healthy donors has shown effectiveness in treating (some) gastrointestinal tract infections (e.g. recurrent *Clostridium difficile* infections [16]) and clinical trials are underway for other diseases indicating the promising future of microbiota transplant as a cost-effective and (hopefully) safer alternative to antibiotics (reviewed in [1,17]).

In addition to genetic, geographical, and social factors that can tailor the composition of the human microbiome, microbial colonization is also challenged by their (in)-ability to occupy certain human body sites [18]. Each body site presents a unique environment and thus limits or favors the growth of certain microorganisms. This generates between body-site variability within the same individuals, which is a hallmark feature of animal (human) microbiome studies. While large-scale metagenome sequencing projects targeting multiple human body sites have already been completed or are underway in industrialized countries (e.g. the Human Microbiome Project [19,20], the European MetaHIT consortium [21], and the White House Microbiome Initiative [22]), the microbiota composition of people living in developing or under-developed countries remains relatively understudied. There are stark differences in the dietary and social habits of people living in, for example, Europe, USA, and Australia compared to people living in, for example, Asia and Africa. Therefore, while microbiome-centric treatments are becoming popular in the industrialized countries, they cannot directly be applied to people living in the developing or under-developed countries [23–25]. Here, we attempt to bridge such knowledge gaps by presenting results of the preliminary sequencing and profiling of gut and oral microbiota in urban Pakistani adults.

With a population of >200 million, Pakistan is the fifth most populous country in the World. It is linguistically, culturally, and geographically diverse and unique in dietary habits. Pakistanis typically consume three meals a day in addition to baked/fried snacks. Although, India and Pakistan share many similarities in diet and culture, Pakistani food composition is considerably different from the Indian food. Meat consumption, for example, is at least three-times higher in an average Pakistani than in an average Indian and is the highest among South Asian countries [26]. Pakistani meals are typically composed of naan/roti bread cooked in oil/ghee and served with vegetable or meat curries enriched in spices and oil (especially in the Punjab region, Figure 1A). Rice consumption is also very high, especially in the Kashmir region, and rice are also typically cooked in oil/ghee along with meat/vegetables. Seafood, especially popular in Sindh, is also preferred fried and in cooked form. Thus, Pakistani way of cooking food that combines oil and spices with vegetables and meat and are served with fried rice or naan/roti (that are also frequently coated with oil/ghee) suggests that Pakistani food is likely a mixture of high-fat and high-starch diet and low in fiber. Fresh fruit intake is pretty low. In fact, 21/31 (67%) people in our study reported no intake of daily fresh fruit (∼50% in a recent survey [27]). Additional sources of carbohydrates are traditional Pakistani desserts such as sweets and puddings that are also loaded with sugar (and sometimes oil such as *mithai* and *jalebi*). Combined with low physical activity (up to 60% in a recent survey [27]) and increasing popularity of “junk food” in urban areas [28], we speculate that Pakistanis typically do not eat or live very healthy. Indeed, a recent study revealed that ∼57% out of 5,491 low-income urban Pakistani individuals had body mass index (BMI) of >25 Kg/m^2^ (i.e. they were overweight/obese) [29]. We also faced significant challenges in recruiting normal BMI participants for the present study and ∼90% of individuals interviewed were either rejected due to being overweight or due to the history of antibiotic consumption (see Discussion). Indeed, Pakistan is number three in the low or middle-income countries in antibiotic consumption (after India and China) [30]. Antibiotic consumption begins from a very early age in Pakistan. A total of 26/31 (84%) recruited study participants reported that they were given antibiotics as children (Table S1). Moreover, ‘self-prescription’ is another common issue in Pakistan [31], where individuals consume antibiotics on their own and sometimes insist doctors to prescribe them drugs.

**Figure 1.**
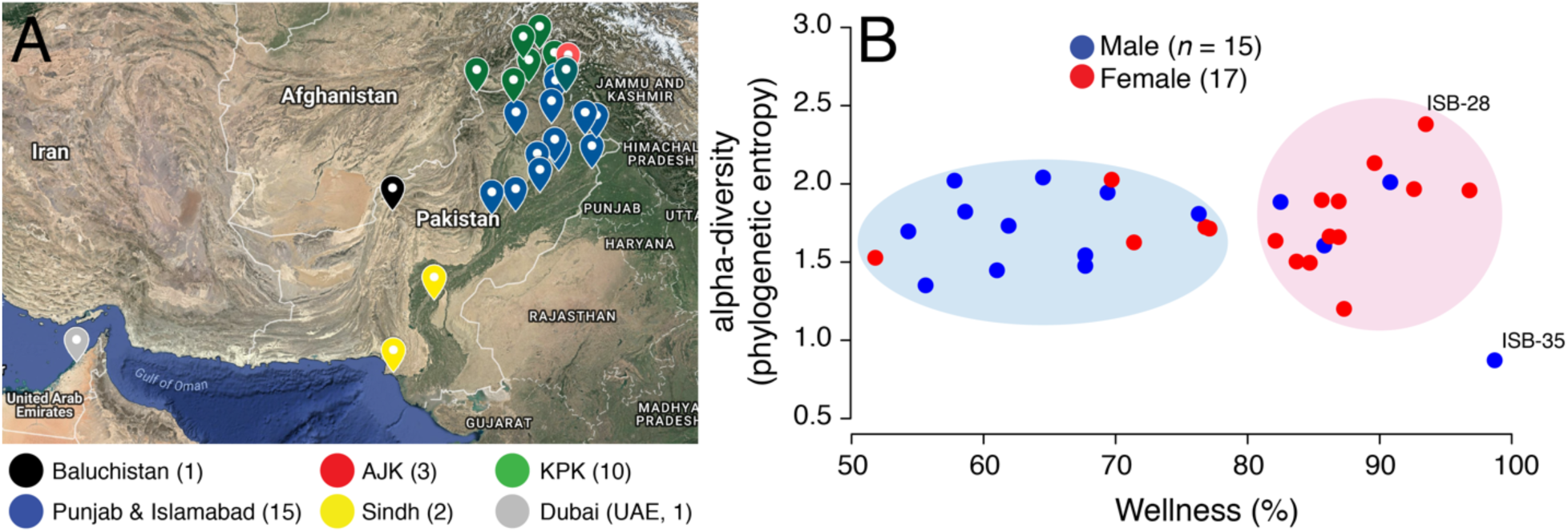
**(A)** Birth cities of study participants are highlighted on the map of Pakistan. Data points are colored according to geographical regions. Islamabad (Federal Capital) was pooled into Punjab for ease of visualization. Map generated online using Google Maps. AJK, Azad Jammu & Kashmir; KPK, Khyber Pakhtunkhwa. **(B)** A scatter-plot displaying relationship between microbiome wellness (%) and alpha-diversity (phylogenetic entropy) for females (red) and males (blue). Wellness percentage match for each sample was retrieved from uBiome Explorer, a commercial product for uBiome users and grant awardees (see Materials and Methods). Individuals with maximum and minimum diversity scores are labeled (see Table S1 for metadata).

Taken together, the dietary habits of Pakistanis of eating fried food enriched in oil and spices, following a relatively sedentary lifestyle (physical inactivity prevalence is ∼60.1% in Pakistan [27]), over-consumption of antibiotics [30], low intake of dietary fiber and fruits, and poor quality of drinking water (and hygienic standards that do not meet international standards) [32] in most regions suggests that Pakistani microbiota would be very different from other populations and perhaps in a permanent state of ‘dysbiosis’ even among carefully selected study participants. Despite these obvious concerns and scientific relevance, to our knowledge, no nation-wide study has hitherto been undertaken to study the composition of the Pakistani microbiome at a large-scale. In collaboration with uBiome, a microbial genomics biotechnology company in San Francisco, CA, we are attempting to characterize the Pakistani microbiome (Pakistan Microbiome Initiative) capturing the full ethnic and geographic diversity of the country. Here, we present results of the pilot phase of the gut and oral microbiome sequencing in young, early-career professionals/students, urban Pakistani adults. Overall, we observed moderate wellness of the Pakistani microbiome, low overall diversity, high levels of probiotics, and strong gender-based differences in the health and wellness of the Pakistani gut. Levels of starch-metabolizing Proteobacteria were abnormally high in Pakistanis, especially males, which could be a marker for dysbiosis and metabolic disorders [33]. Our effort highlights key areas we need to focus on in the subsequent phases of the Pakistani Microbiome Initiative, is the first step towards generating a country-wide understanding of the impact of the Pakistani diet and lifestyle on Pakistani gut microbiota, and yields actionable insights to hopefully improve diet and wellness practices in Pakistan.

## MATERIALS AND METHODS

### Participant Identification and Screening

We recruited study participants through phone and in-person interviews conducted at COMSATS University Islamabad (project sampling site). The major exclusion criteria were body mass index (BMI) either <18 or >25 Kg/m^2^, prior history of colon cancer, pregnant or lactating women, or women with irregular menstrual cycles (i.e. less than 21 or more than 35 days apart). We attempted to capture maximum gender, geographic, and ethnic diversity of the country. Our shortlisted study participants were therefore 32 individuals (17 females and 15 males, mean age = 23.41, SD = ± 4.83 years, Tables 1 and S1 for metadata) who donated a total of 61 stool and saliva samples and were born in 25 major cities (Figure 1A) representing seven major geographic regions (Punjab, Khyber Pakhtunkhwa (KPK), Baluchistan, Azad Jammu and Kashmir (AJK), Federal Capital Islamabad, Sindh, and United Arab Emirates (UAE) and six major ethnicities (Baloch, Kashmiri, Pashtun, Punjabi, Saraiki, and Sindhi) (Tables 1 and S1). All individuals had ‘normal’ BMI values ranging from 18.1 Kg/m^2^ to 24.9 Kg/m^2^ (mean = 22.0, SD = ± 2.14 Kg/m^2^). Six individuals reported antibiotic intake in the three months prior to sampling, four individuals reported acute or chronic diarrhea in the two months prior to sampling, one individual had history of inflammatory bowel disease (IBD), five individuals had been diagnosed in the past with a medical condition (chronic liver disease followed by liver transplant, rheumatoid arthritis, hepatitis A, and tuberculosis), while one individual experienced constipation in the two-month period prior to sampling. Together they constituted the “cases” group comprising of 13 individuals (25 samples) that could be contrasted to the relatively normal or “control” group of 19 individuals (36 samples) who did not report any ailments (Table 1). In addition, one individual reported international travel outside Pakistan (United Arab Emirates) in the six-month period prior to sampling. The Ethics Review Board at COMSATS University Islamabad approved the study. All participants provided written informed consent to participate. Participant data was analyzed in aggregate and anonymously.

**Table 1.** Characteristics of shortlisted study participants. A total of 32 individuals (17 females and 15 males) donated 61 samples (32 gut and 29 oral). KPK, Khyber Pakhtunkhwa; AJK, Azad Jammu and Kashmir; UAE, United Arab Emirates (see Table S1 for detailed metadata). BMI, body mass index.

| ID | Samples Donated | Province of Birth | Ethnicity | Gender | Age | BMI | Class | Disclosure |
| --- | --- | --- | --- | --- | --- | --- | --- | --- |
| ISB-01 | Stool & Saliva | Punjab | Punjabi | Female | 24 | 24.6 | Cases | History of inflammatory bowel disease and tuberculosis |
| ISB-04 | Stool & Saliva | KPK | Pashtun | Male | 25 | 22.4 | Control |  |
| ISB-05 | Stool & Saliva | KPK | Pashtun | Male | 26 | 24 | Control |  |
| ISB-06 | Stool & Saliva | Punjab | Punjabi | Female | 25 | 24.2 | Cases | Antibiotic intake in the past three months |
| ISB-07 | Stool & Saliva | AJK | Kashmiri | Female | 20 | 20.6 | Control |  |
| ISB-09 | Stool & Saliva | Punjab | Punjabi | Female | 18 | 18.7 | Control |  |
| ISB-10 | Stool & Saliva | Punjab | Punjabi | Female | 26 | 23.5 | Control |  |
| ISB-11 | Stool & Saliva | Punjab | Punjabi | Male | 30 | 23 | Control |  |
| ISB-12 | Stool & Saliva | AJK | Kashmiri | Female | 24 | 21.5 | Control |  |
| ISB-13 | Only stool | Punjab | Punjabi | Female | 19 | 21.1 | Control |  |
| ISB-14 | Stool & Saliva | UAE | Pashtun | Female | 23 | 21.9 | Control | UAE travel in the past six months |
| ISB-15 | Stool & Saliva | KPK | Pashtun | Female | 26 | 24.2 | Control |  |
| ISB-17 | Only stool | Sindh | Sindhi | Female | 19 | 24.2 | Cases | Diarrhea in the past two months |
| ISB-18 | Stool & Saliva | Punjab | Punjabi | Female | 21 | 22.3 | Control |  |
| ISB-19 | Stool & Saliva | Federal Capital | Punjabi | Female | 18 | 21.6 | Cases | Unnamed medical condition |
| ISB-20 | Stool & Saliva | Punjab | Punjabi | Female | 24 | 21.4 | Cases | Antibiotic intake in the past three months. Diagnosed with Rheumatoid arthritis and Hepatitis A. |
| ISB-21 | Stool & Saliva | Sindh | Punjabi | Female | 24 | 19.2 | Cases | Constipation at the time of sampling |
| ISB-24 | Stool & Saliva | Punjab | Punjabi | Male | 20 | 18.5 | Cases | Antibiotic intake in the past three months. History of Malaria. |
| ISB-27 | Stool & Saliva | KPK | Pashtun | Male | 19 | 23.5 | Control |  |
| ISB-28 | Stool & Saliva | Punjab | Punjabi | Female | 24 | 23.8 | Control |  |
| ISB-29 | Stool & Saliva | KPK | Pashtun | Male | 18 | 20.9 | Control |  |
| ISB-30 | Stool & Saliva | KPK | Pashtun | Male | 18 | 20.9 | Cases | Antibiotic intake in the past three months |
| ISB-31 | Stool & Saliva | KPK | Pashtun | Male | 18 | 19.5 | Cases | Antibiotic intake in the past three months |
| ISB-32 | Stool & Saliva | KPK | Pashtun | Male | 23 | 24.7 | Control |  |
| ISB-33 | Stool & Saliva | Punjab | Saraiki | Female | 24 | 18.1 | Control |  |
| ISB-34 | Stool & Saliva | Punjab | Saraiki | Female | 35 | 23.9 | Control |  |
| ISB-35 | Stool & Saliva | Punjab | Saraiki | Male | 40 | 24.9 | Cases | Chronic liver disease, liver transplant |
| ISB-36 | Stool & Saliva | Baluchistan | Baloch | Male | 26 | 20.1 | Cases | Diarrhea in the past two months |
| ISB-37 | Stool & Saliva | AJK | Kashmiri | Male | 24 | 21 | Cases | Antibiotic intake in the past three months & Diarrhea in the past two months |
| ISB-38 | Stool & Saliva | KPK | Pashtun | Male | 23 | 18.2 | Cases | Diarrhea in the past two months |
| ISB-39 | Stool & Saliva | Punjab | Punjabi | Male | 22 | 24.8 | Control |  |
| ISB-40 | Only stool | KPK | Punjabi | Male | 23 | 24.1 | Control |  |

### Sample Collection

Stool and saliva samples were self-collected by participants at home in uBiome commercial kits. These kits follow the protocols outlined by the NIH Human Microbiome Project [34]. Participants were instructed to use sterile swabs to transfer small amounts of fecal material and saliva into vials containing lysis and stabilization buffer for DNA storage at room temperature.

### 16S Ribosomal RNA (rRNA) Gene Sequencing, DNA Extraction, and PCR Amplification

These steps were performed in the CLIA-compliant (Clinical Laboratory Improvement Amendments) and CAP-accredited (College of American Pathologists) uBiome laboratory in San Francisco, CA. Samples were lysed using mechanical bead-beating [35]. DNA was extracted and purified by a liquid-handling robot in a class 100 clean room using the method described in [36]. The V4 region of 16S rRNA gene was PCR amplified using universal forward and reverse primers (515F: GTGCCAGCMGCCGCGGTAA and 806R: GGACTACHVGGGTWTCTAAT). Primers also contained Illumina tags and barcodes with unique combination of forward and reverse indexes to allow multiplexing. Pooled PCR products were column-purified and selected through microfluidic DNA fractionation based on size [37]. Real-time qPCR quantified consolidated libraries using the Kapa Bio-Rad iCycler qPCR kit on a BioRad MyiQ prior to sequencing.

### DNA Sequencing and Quality Control

16S amplicons from each sample were individually barcoded and sequenced in multiplex in the NextSeq 500 platform in a 150bp paired-end modality. Raw data from the sequencer was first demultiplexed, and the forward and reverse reads obtained in each of the four lanes per sample were filtered using the following criteria: (i) Both forward and reverse reads in a pair must have an average Q-score > 30, and (ii) primers, and any leading random nucleotides (used to increase diversity of the library being sequenced) were trimmed, and forward reads were capped at 125bp and reverse reads were capped at 124bp. Forward and reverse reads of each pair were appended, and those sequences that contained more than 8 consecutive same nucleotide repeats were discarded. Remaining sequences were clustered using a distance of 1 nucleotide using the Swarm algorithm [38] and the most abundant sequence per cluster was considered the representative of the cluster and assigned a count corresponding to the sum of sequences that composed the cluster. A chimera removal using these centroid representative sequences was performed using the VSEARCH uchime_denovo algorithm [39]. Singletons that remained after chimera removal were also discarded.

### Taxonomy Assignment

Both forward and reverse reads that matched with >77% sequence identity to the same sequence in version 123 of the SILVA database retrieved from https://www.arb-silva.de [40] were assumed to be 16S sequences. The most abundant forward-reverse read pair per Swarm cluster was assigned taxonomic annotation according to the following thresholds: >97% identity (species), >95% (family), >85% (order), >80% (class), >77% (phylum). In total, we assigned taxonomy to ∼5.5 million read-pairs (total = 5,580,520, 174,391/sample, minimum = 7,847, maximum = 353,529) in 32 gut samples and to ∼4.3 million sequences (total = 4,380,874, 151,064/sample, minimum = 5,805, maximum = 386,012) in 29 oral samples. Two samples that were <10,000 read count threshold were dropped (as in [41]) for downstream diversity and comparative analyses. These included stool sample donated by ISB-35 and saliva sample donated by ISB-11 (Tables 1 and S1). ISB-35 had recently undergone liver transplant and was taking immunosuppressants that likely reduced his gut microbiota diversity. ISB-11, however, appeared healthy, as per our questionnaire (Table S1) but had abnormally low-counts of oral bacteria relative to other samples. To ensure consistency in target taxa detection even at low abundances, this sample was also removed in downstream analyses. Remaining samples (*n* = 59, >10,000 taxa) were rarefied to minimum library sizes (gut = 14,574/samples, oral = 11,878/sample) as there was still >10X difference between the library sizes of few small samples versus the maximum library size necessitating rarefaction [42].

### Bioinformatics Analysis

MicrobiomeAnalyst [43] was used to perform downstream analyses. A taxonomy abundance table containing features (phyla, genera, and species) and their abundance scores (detected raw counts at phyla, genera, and species levels) for all samples was provided as input along with sample metadata (Tables 1 and S1). Low-count and low-variance features were removed keeping only features with >4 count in at least 20% samples. Raw counts were rescaled using the method of total sum scaling. Weighted UniFrac distance was used to plot sample dissimilarity on the two main principal coordinate axes. Statistical significance of dissimilarity between study groups (e.g. gender, control vs. cases) was evaluated by the Analysis of Similarities (ANOSIM) method implemented in MicrobiomeAnalyst. Within-sample diversity (i.e. alpha-diversity) was evaluated by abundance-weighted phylogenetic entropy index [44], which is linearly dependent on the branch lengths in the phylogenetic tree and increases with distinctiveness of the sample [45], along with traditional non-phylogenetic alpha-diversity measures (observed, Chao1, and Shannon’s diversity). Linear Discriminant Analysis Effect Size (LEfSe) algorithm was implemented for biomarker discovery [46]. The algorithm first computes differential abundances of features across study groups followed by LDA to determine their effect size or relevance. Differential abundance of taxa was evaluated for key metadata collected from participants including gender, geography, ethnicity, dietary and social habits, and medial health questions (Table S1). Statistical significance was evaluated either by two-tailed Wilcoxon rank sum test (2 samples) or Kruskal-Wallis (>2 samples), where appropriate (*P* < 0.05, *FDR* < 0.05). Abundance correlations were determined by Kendall rank correlation coefficient (τ) and visualized by heat-maps.

### uBiome Citizen Science Initiative

Relative abundances of major phyla and genera detected in Pakistani gut and oral samples were compared against corresponding abundances in the gut and oral samples of self-reported healthy individuals participating in the ongoing uBiome Citizen Science initiative (e.g. [41]). These data are available from the uBiome Explorer website to all commercial users and to uBiome academic grant awardees for research purposes. The uBiome healthy individuals completed a detailed questionnaire about 42 different medical conditions covering infectious diseases, metabolic disorders, chronic health issues, and mental health disorders and had never been diagnosed with abnormal blood glucose levels, diabetes, or digestive-tract related disorders or any other medical condition (see [41]). Informed consent was obtained from all participants and these individuals consented to share their results for research purposes. This study was performed under a Human Subjects Protocol provided by an IRB (E&I Review Services, IRB Study #13044, 05/10/2013).

## RESULTS

### Pakistani Gut Microbiome is Moderately Healthy Relative to Western Populations

We compared the gut microbiota composition of Pakistani samples against the uBiome dataset. A wellness match percentage, defined by uBiome Explorer, is the overlap in the microbiome composition between the samples of interest (Pakistani samples) and selected samples from individuals who report no ailments (see [41] for an example study on the uBiome healthy cohort). Thus, it can be a useful proxy to evaluate whether the individual’s microbiota resembles that of microbiota in healthy individuals or not (see Discussion for limitations of using this index). Wellness percentage in Pakistani individuals ranged from 51.80% to 98.70% (mean = 76.73%, SD = ±13.55%) (Figure 1B) indicating overall moderate health and wellness of the Pakistani gut microbiome relative to worldwide (Western) populations. Interestingly, wellness percentage was significantly higher in females versus males (mean = 82.51% vs. 70.17%, *P* = 0.01, two-tailed Wilcoxon rank sum test) suggesting that Pakistani women on average harbored a gut microbiome profile that was relatively more similar to the gut microbiome profiles in healthy individuals.

In fact, Pakistani samples segregated into two major clouds when plotted against wellness match and alpha-diversity (i.e. the number and distribution of taxa within a sample [44]), as measured by the phylogenetic entropy index [45] (Figure 1B). The right cloud included samples with wellness match >80% and included 12/17 (71%) females. In comparison, the left cloud exhibited wellness match scores between 50-80% and predominantly included males (11/15, 73%). Phylogenetic entropy ranged from 0.873 (lowest, male) to 2.381 (highest, female) with a mean of 1.727, SD = ±0.29. While, numerically females had higher mean alpha-diversity compared to males (1.76 vs. 1.68), the difference was statistically insignificant (*P* = 0.62, two-tailed Wilcoxon rank sum test). The male individual with the lowest diversity (ISB-35) had recently undergone liver transplant after chronic liver infection (Table S1). This individual was taking immunosuppressants that significantly reduced his gut microbiome diversity (removed from subsequent analyses). In comparison, female with maximum gut diversity (ISB-28) was interestingly the only one who reported no intake of “naan” (Table S1), a typical Pakistani/Indian flatbread made from wheat flour and served usually with ghee/oil. Naan/roti are rich in carbohydrates and a mandatory part of almost every Pakistani meal, especially in Punjab. This individual was therefore likely on “low-carb” diet relative to all other Pakistanis who consume high-carb and high-fat, presumably, unhealthy meals (see Discussion). Other than gender, however, wellness scores and diversity of samples did not differ significantly either by geography or ethnicity (Figure S1). Finally, there was no significant difference in either the wellness match or diversity scores between control and cases (*P* = 0.76 and *P* = 0.62, two-tailed Wilcoxon rank sum test, Figure S1).

### Pakistani Gut Microbiome Appears More Similar to the Western Microbiome Relative to its Geographical Neighbor

After removal of ISB-35 and quality-control and filtering (see Materials and Methods), we determined that four major or core phyla, Firmicutes, Bacteroidetes, Proteobacteria, and Actinobacteria, dominated the Pakistani gut microbiome community comprising, on average, ∼96% of total bacteria load (Figure 2A, Tables 2 and S2). There were, however, large fluctuations in the individual proportions of each of these core phyla across individuals (Figure 2A), which is not surprising since individuals typically follow a continuum rather than clustering into discrete groups when compared across a particular body site [47,48]. Interestingly, the gut microbiome profile in Pakistan is more similar to the uBiome profile rather than its geographical and cultural neighbor (India). For example, in a recent study conducted in Western India, these four phyla also represented ∼95% of the gut bacterial community, albeit with starkly different compositions [23]. The Indian gut microbiota was heavily skewed towards Bacteroidetes (∼71% of total bacteria) whereas Pakistani gut microbiota appears skewed towards Firmicutes (46.2%, Bacteroidetes = 24.5%, Figure 2B, Table 2) and, in addition, also has a very high Proteobacteria load (15.86% vs. 3.8% in [23]). These differences are likely due to dietary differences between the two countries, especially, since meat consumption is very high in Pakistan relative to India [26] and Indians typically prefer a plant-based diet. In this regard, Pakistani microbiome shares more similarities to the uBiome healthy cohort (Figure 2B). For example, the gut microbiomes of uBiome healthy individuals are also skewed towards Firmicutes (60.1%) whereas Bacteroidetes constitute roughly ∼30% of the total microbial community (Table 2). Compared to uBiome healthy individuals, however, both Pakistani males and females harbored lower mean proportions of Firmicutes (40.1% and 51.9% vs. 60.1% in uBiome) and Bacteroidetes (21.18% and 27.24% vs. 31.06%) but higher mean proportions of Proteobacteria (25.06% and 8.29% vs. 3.83%) and Actinobacteria (9.31% and 7.99% vs. 4.1%) (Figure 2B, Table 2). While, mean proportions of all four phyla were different from uBiome healthy individuals for both Pakistani men and women, women appeared to more closely resemble the numbers in healthy populations (Table 2). There were no major differences in the composition of these four core phyla between control and cases (Figure S2).

**Table 2.** Mean relative abundance of major phyla (detected in 100% gut samples) and their comparison to the uBiome dataset (see Table S2 for complete list).

| Phylum | Mean relative abundance (all) |  |  | Mean relative abundance (males) |  |  | Mean relative abundance (females) |  |  | uBiome<sup>a</sup> |
| --- | --- | --- | --- | --- | --- | --- | --- | --- | --- | --- |
|  | *min* | *mean* | *max* | *min* | *mean* | *max* | *min* | *mean* | *max* |  |
| Firmicutes | 21.80 | 46.62 | 76.36 | 23.73 | 40.14 | 58.14 | 21.80 | 51.96 | 76.36 | 60.1 |
| Bacteroidetes | 11.79 | 24.50 | 51.45 | 13.00 | 21.18 | 31.06 | 11.79 | 27.24 | 51.45 | 31.06 |
| Proteobacteria | 2.07 | 15.87 | 41.85 | 5.53 | 25.07 | 41.85 | 2.07 | 8.29 | 18.53 | 3.83 |
| Actinobacteria | 0.68 | 8.59 | 37.25 | 0.68 | 9.31 | 37.25 | 1.69 | 8.00 | 16.12 | 4.1 |

<sup>a</sup>Numbers as reported on uBiome Explorer in July 2018. Numbers are subject to change with inclusion of more healthy participants to the ongoing uBiome citizen science initiative (e.g. [41]).

**Figure 2.**
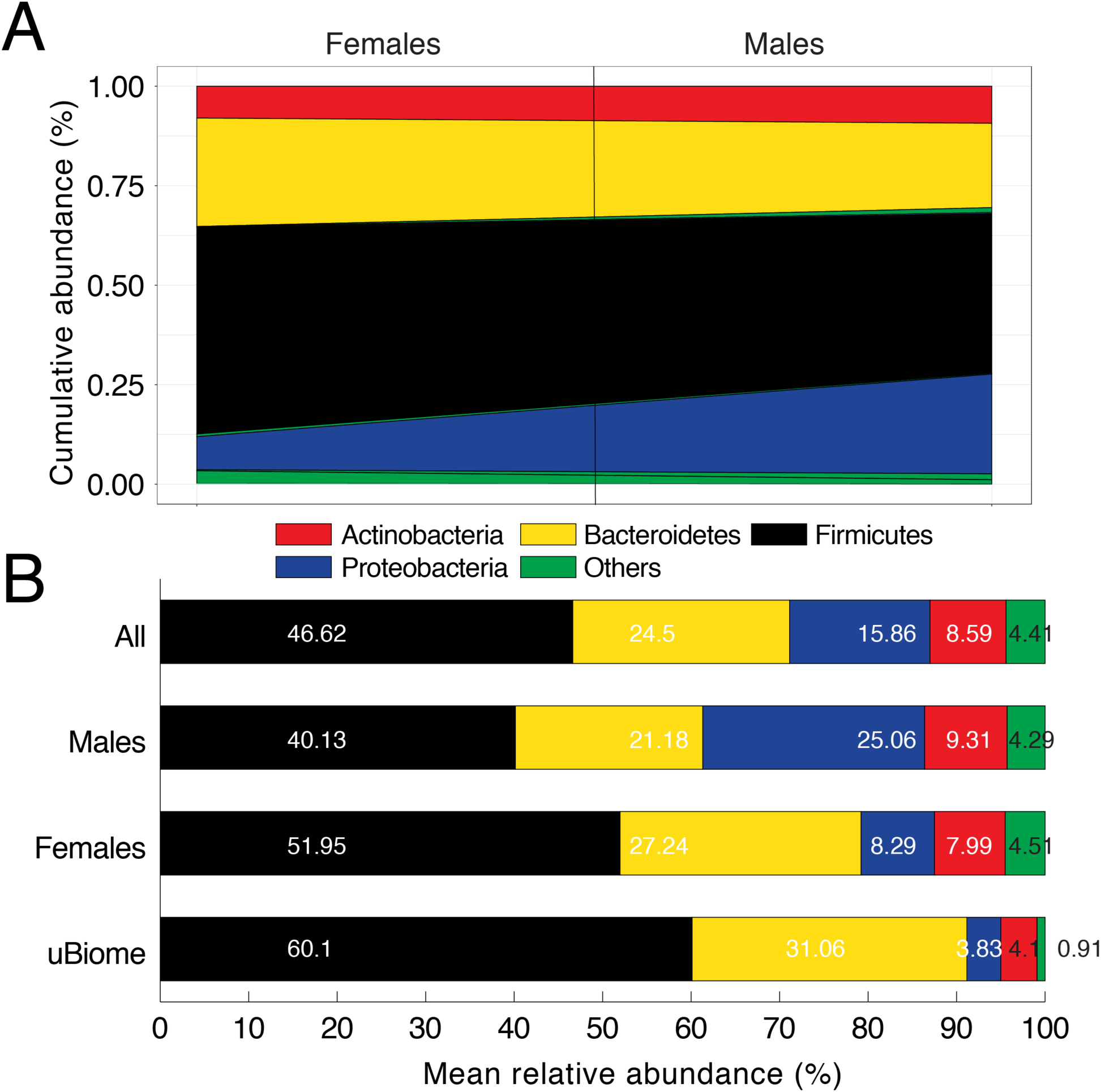
**(A)** Area diagram displaying composition of microbial phyla detected in Pakistani gut samples. Black line separates male and female samples. *Others* category includes Tenericutes, Lentisphaerae, Euryarchaeota, Elusimicrobia, Verrucomicrobia, Fibrobacteres, Spirochaetes, and Synergistetes (Table S2 for actual numbers for all detected phyla). **(B)** Comparison of mean relative abundance of microbial phyla for Pakistani males and females versus uBiome dataset. Numbers on bars indicate actual percentages.

### Abnormally High Levels of Proteobacteria in Pakistani (Male) Samples Could Be of Concern

High load of Proteobacteria in Pakistani individuals relative to both Western Indians and uBiome datasets (mean relative abundance = 15.8% vs. 3.8%) could be of concern, especially since it has been associated with various metabolic disorders and is a marker of dysbiosis (reviewed in [33]). Proteobacteria load was especially very high in Pakistani men ranging from 5.53% to 41.84% (mean = 25.06%) compared to 2.06% to 18.53% (mean = 8.29%) in women (Table 2). In other words, even the minimum Proteobacteria load detected in any Pakistani male (5.53%) was still higher than the average Proteobacteria load expected in healthy individuals. A differential abundance analysis followed by linear discriminant analysis (LDA) confirmed these gender-wise differences. Proteobacteria, Actinobacteria, Spirochaetes, Elusimicrobioa, and Fibrobacteres characterized male samples (LDA >3) whereas Firmicutes, Bacteroidetes, Tenericutes, Lentisphaerae, Verrumicrobia, Euryarchaeota, and Synergistetes characterized female samples (Figure 3A). Relative abundances of Proteobacteria, Spirochaetes and Elusimicrobia were significantly higher in males relative to females (*FDR* < 0.05) whereas women had a relatively higher load of Firmicutes to males, albeit with marginal significance (*P* = 0.03, *FDR* = 0.1) (Figure 3B, Table 3). Abundance of Proteobacteria was moderately correlated with Spirochaetes (Kendall’s τ = 0.42) and negatively correlated with Firmicutes (τ = −0.49) (Figure 3C, see Table S3 for correlation scores). Previously, Proteobacteria load of >13% has been associated with metabolic disorders and inflammation and cancer (reviewed in [33]). Thus, enrichment of Proteobacteria in Pakistani males could be of concern since the majority of recruited participants self-reported to be in good overall health and had normal BMI (Tables 1 and S1). Time series monitoring is therefore warranted to test whether proteobacteria are indeed chronically enriched in the Pakistani gut or not (see Discussion). Two additional metadata variables were also associated with differential abundance of phyla including stress levels in the one-week prior to sampling and income group. In stress, Spirochaetes were significantly depleted in individuals who experienced any level of stress (extreme, moderate, slight) versus no stress (Figure S3, *FDR* < 0.05) whereas phyla with significant differential abundance across income levels (Lentisphaerae and Verrumicrobia) did not rise or decline in a manner consistent with income growth (Table S4). Other than these three variables (gender, stress, income), however, none of the additional key metadata variables (including questions regarding ethnicity, geography, marital status, socioeconomic status, dietary preferences, Table S1) were associated with differential abundance of phyla in the Pakistani gut.

**Table 3.** Differential abundance analysis of microbial phyla by gender for Pakistani gut samples. Statistically significant differences (*FDR* < 0.05) are highlighted. LDA scores are mentioned.

| Phylum | *P*-value | *FDR* | Female | Male | LDA score |
| --- | --- | --- | --- | --- | --- |
| Proteobacteria | 0.00011795 | 0.0014155 | 829196.23 | 2506616.48 | 5.92 |
| Elusimicrobia | 0.0016788 | 0.010073 | 3269.32 | 122380.36 | 4.77 |
| Spirochaetes | 0.0085595 | 0.034238 | 24297.90 | 147473.98 | 4.79 |
| Firmicutes | 0.035396 | 0.10619 | 5195997.71 | 4013850.50 | -5.77 |
| Verrucomicrobia | 0.33017 | 0.7924 | 22279.81 | 2205.49 | -4 |
| Bacteroidetes | 0.4998 | 0.88298 | 2724150.18 | 2118106.61 | -5.48 |
| Synergistetes | 0.73776 | 0.88298 | 2583.17 | 1764.39 | -2.61 |
| Tenericutes | 0.77991 | 0.88298 | 316841.43 | 110617.73 | -5.01 |
| Fibrobacteres | 0.78056 | 0.88298 | 4601.26 | 16957.79 | 3.79 |
| Euryarchaeota | 0.79433 | 0.88298 | 11382.07 | 5244.17 | -3.49 |
| Lentisphaerae | 0.8094 | 0.88298 | 65870.73 | 23329.22 | -4.33 |
| Actinobacteria | 0.93672 | 0.93672 | 799530.19 | 931453.27 | 4.82 |

**Figure 3.**
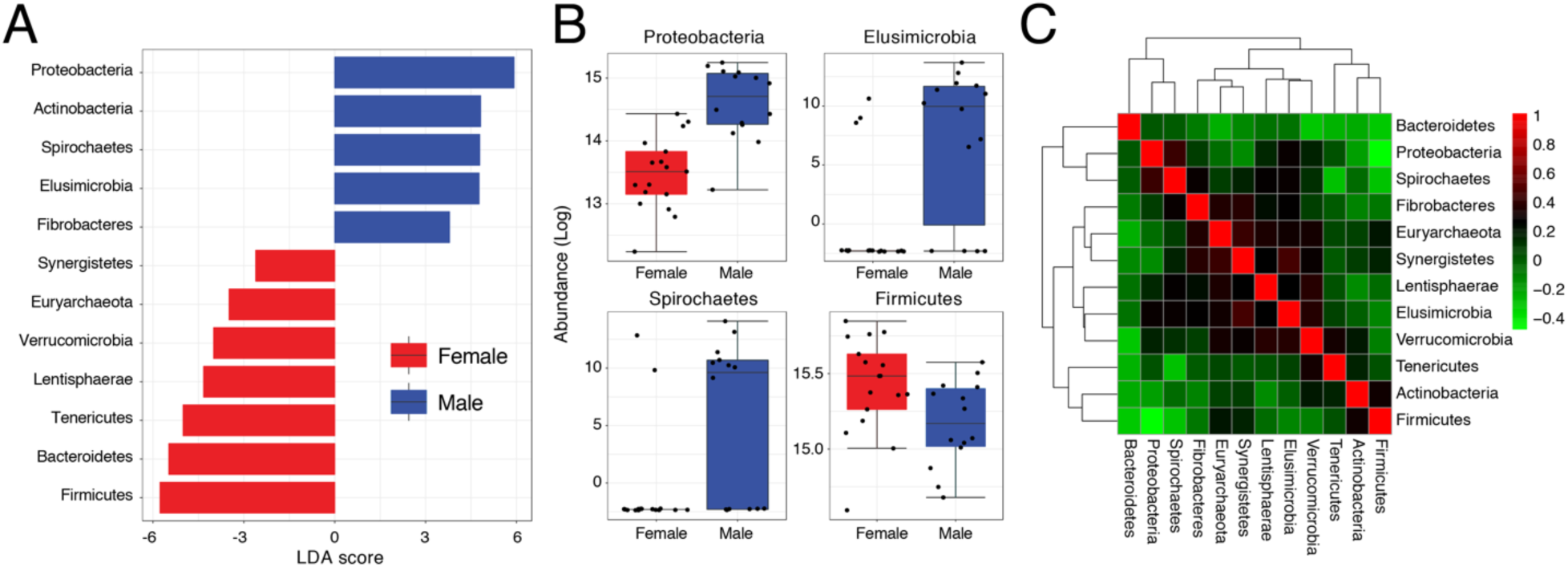
**(A)** Bar graph showing LDA scores of significant features (phyla) characterizing Pakistani male and female gut samples. **(B)** Boxplots comparing phyla with significant differential abundance in males and females (Wilcoxon rank-sum test, *FDR* < 0.05). **(C)** A heat-map visualizing how phyla abundances in the Pakistani gut correlate to each other (Kendall’s τ).

### Pakistani Gut Microbiota is Tilted Towards Weight Gain

Firmicutes and Bacteroidetes are the two most prominent members in the gut of human populations worldwide [49]. Firmicutes are efficient in fat digestion and their increased representations in the human gut have been linked to obesity (see [50,51] for contradictory evidence). In contrast, Bacteroidetes do not digest fat very well and are thought to help protect against obesity and weight gain [52,53]. Bacteroidetes are typically under-represented in the gut of Europeans and North Americans (similar to Pakistani microbiome, Table 2), however, this is likely due to diet and lifestyle than genetics [54,55]. Firmicutes to Bacteroidetes ratio has been associated with tendencies towards weight gain and loss. Microbiomes skewed in favor of Firmicutes have been correlated with weight gain and obese body types while microbiomes tilted in favor of Bacteroidetes correlate with weight loss and lean body types [56]. Firmicutes are roughly two times the population of Bacteroidetes in the uBiome healthy dataset (2:1, Table 2). In turn, the mean Firmicutes to Bacteroidetes ratio in Pakistani samples is 2.39 (males = 2.07, females = 2.65, Figure 4). In fact, 26/31 (84%) of sampled individuals had Firmicutes to Bacteroidetes ratio of >1 (shaded areas in Figure 4). Despite the fact that the Pakistani gut microbiome is skewed towards Firmicutes, both Pakistani men and women had relatively fewer Firmicutes and Bacteroidetes compared to uBiome dataset (Figure 4B). Only 5/17 (29%) and 6/17 (35%) women had comparable or greater number of Firmicutes and Bacteroidetes relative to uBiome individuals whereas only 1/14 men had comparable or greater number of Bacteroidetes relative to uBiome individuals (shaded area in Figure 4B). This suggests that despite relatively lower proportions of these two key phyla in the Pakistani gut, there is much greater imbalance and, in general, the Pakistani gut microbiome is heavily skewed towards Firmicutes (i.e. towards weight gain and obesity according to some studies [56]). A low-carbohydrate diet, meat without antibiotics, and fruits enriched in polyphenols (apples, pears, berries) are therefore recommended to increase the number of Bacteroidetes in the Pakistani gut.

**Figure 4.**
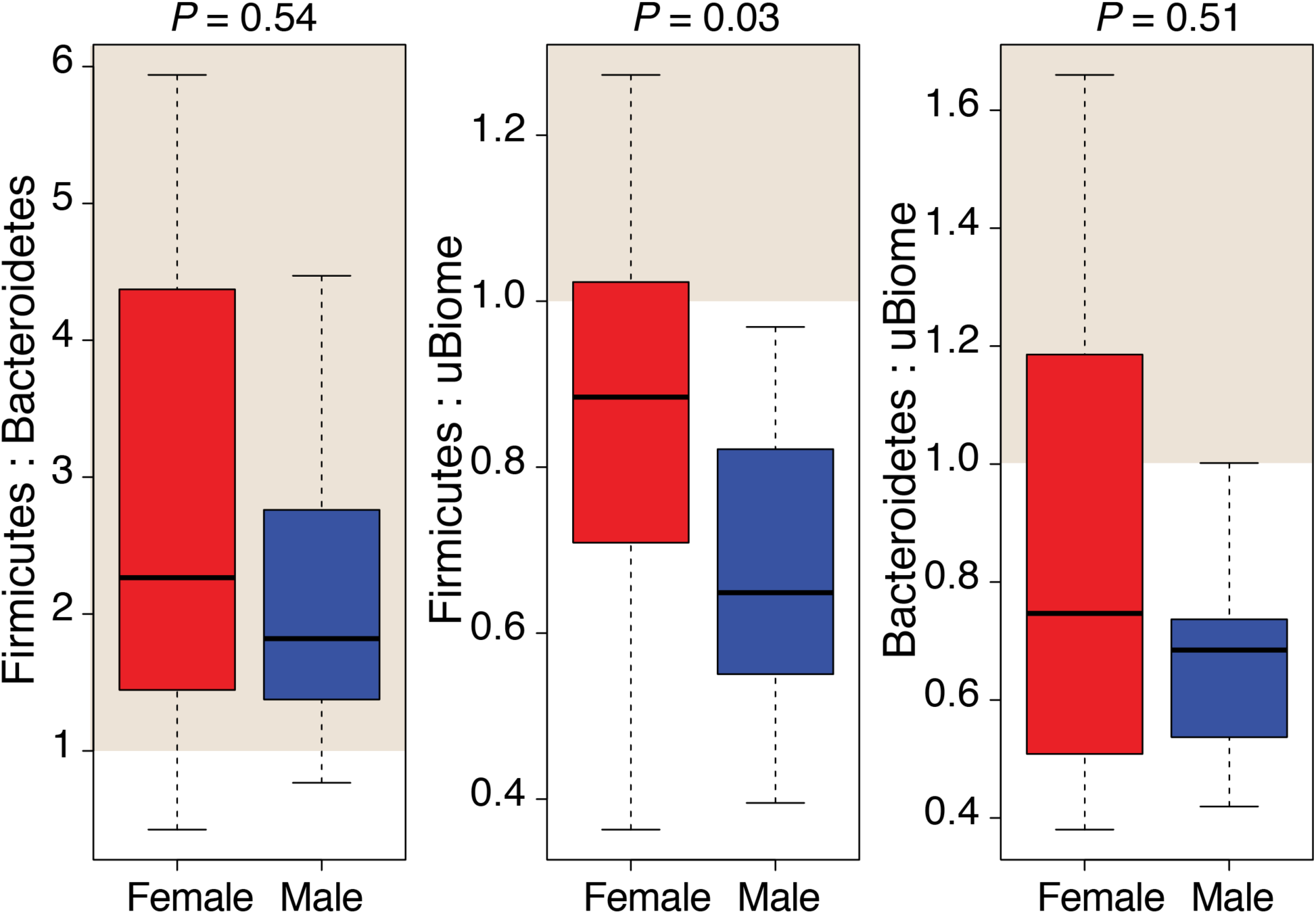
Firmicutes to Bacteroidetes ratio in Pakistani male and female gut samples and their comparison to uBiome dataset. Shaded areas indicate samples where ratio exceeds 1. Statistical significance evaluated by two-tailed Wilcoxon rank sum test (*P* < 0.05).

### Pakistani Men Have Abnormal Levels of Starch-Metabolizing Bacteria

Interestingly, the most abundant genus in Pakistani gut was *Succinivibrio* (Proteobacteria) that was detected in 19/32 samples with a mean relative abundance of 12.21% (Table S5). However, it was especially more prevalent and abundant in males (14/14 samples and mean relative abundance = 23.46%) than females (5/17 samples and mean = 2.94%) (Table S5). Beta-diversity using weighted UniFrac distance reasonably separated male and female samples (54.8% variability explained by two principal coordinates) and confirmed enrichment of *Succinivibrio* in male samples (Figure 5A). *Succinivibrio*, along with *Lachnospira* (Firmicutes), *Elusimicrobium* (Elusimicrobia), and *Phascolarctobacterium* (Firmicutes) had significant differential abundance in males and females (Figure 5B, *FDR* < 0.05). Except *Lachnospira*, rest were significantly over-represented in males (Figure 5B). In Pakistani samples, *Succinivibrio* was positively correlated with *Phascolarctobacter* (τ = 0.55, Figure S4, Table S6). *Succinivibrio* are typically over-represented in cattle switched from high-fiber to high-starch diet [57] and are integral members of the honey bee gut microbiota, which also relies on starch [58]. Thus, they are likely associated with starch metabolism [59]. Pakistani diet is indeed enriched in carbohydrates due to daily, multiple times, intake of rice and naan/roti. The high abundance of *Succinivibrio* in Pakistani gut thus could be expected. However, their very high abundance and prevalence in Pakistani males are especially noteworthy. We speculate that it could be because women have fewer calorie needs than men, are more diet conscious (especially before wedding in Pakistan), and are more dissatisfied with their weight and body shape [60]. Thus, they are likely to consume relatively smaller portions of high-starch Pakistani diet relative to men. An LDA determined 25 most significant features (genera) and their effect sizes (relevance) in categorizing males and females (Figure 5C). Along with *Succinivibrio, Elusimicrobium, Thalassospira, Phascolarctobacterium*, and *Varibaculum* characterized male samples with LDA scores of >|3| whereas *Faecalibacterium, Subdoligranulum, Collinsella, Alistipes, Lactobacillus, Clostridium, Dorea* (and several others) characterized female samples with LDA score >|3| (Table S7 for all features).

**Figure 5.**
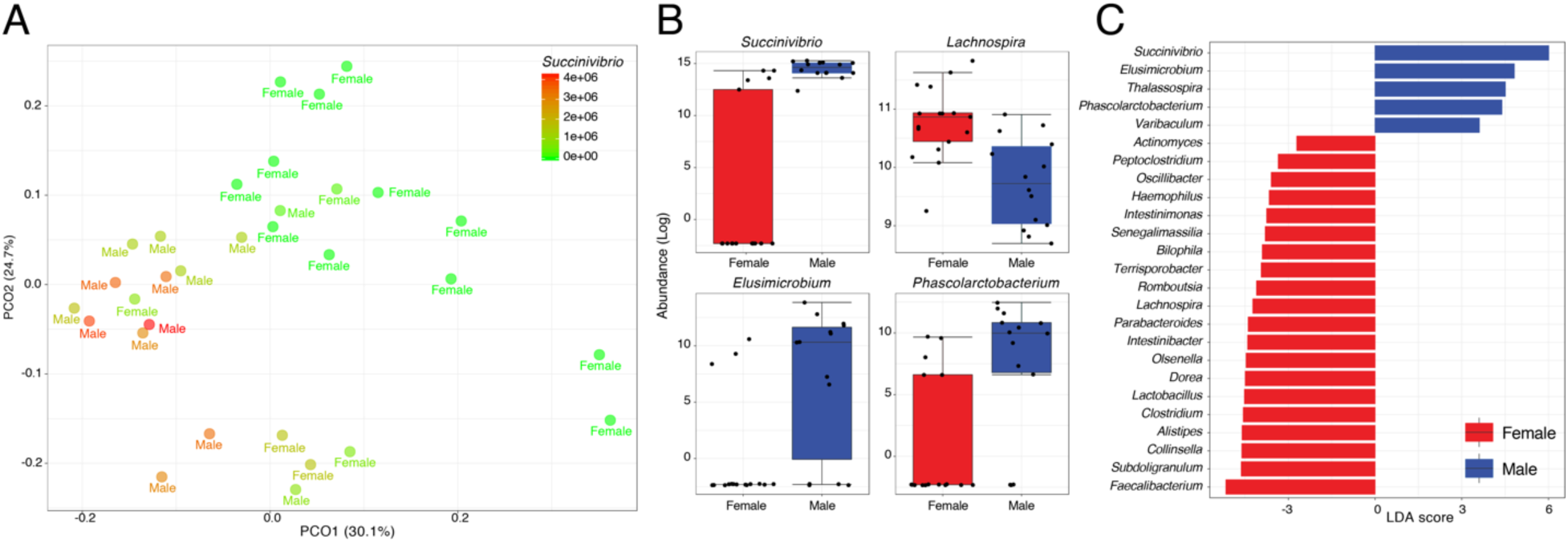
**(A)** A two-dimensional PCoA plot evaluates dissimilarity (Weighted UniFrac) between male and female samples. Samples are colored by *Succinivibrio* abundance. **(B)** Boxplots comparing genera with significant differential abundance in males and females (Wilcoxon rank-sum test, *FDR* < 0.05). **(C)** Bar graph showing LDA scores of top 25 significant features (genera).

### A Coarse-Grained Snapshot of the Pakistani Gut Microbiome

In the West Indian gut microbiome study, *Faecalibacterium, Bacteroides, Roseburia, Alloprevotella,* and *Prevotella* genera comprised 80% of total bacteria in the gut [23]. The microbiome was largely dominated by *Prevotella*, which has been associated with fiber-rich diet [54,61]. Again, Pakistani gut microbiome at genus level is significantly different from the Indian study. The Pakistani gut is dominated by *Succinivibrio* (Proteobacteria), which may, however, be a marker of dysbiosis [33], and *Prevotella* is present at very low levels (mean relative abundance = 0.81% in 20/31 samples, Table S5). In turn, the uBiome dataset indicates dominance of *Faecalibacterium, Bacteroides, Roseburia*, and *Blautia,* comprising >50% of total bacteria (Table 4). In Pakistani samples, 18 different genera (that were detected in all 31 samples) collectively comprised ∼50% of the total gut microbial community (excluding *Succinivibrio*). These 18 genera included notable inhabitants of the human gut including *Faecalibacterium, Bacteroides, Roseburia, Blautia*, and probiotics such as *Bifidobacterium* (Table 4). Interestingly all five genera were under-represented in Pakistani samples relative to uBiome dataset and once again Pakistani women had relatively higher mean abundances compared to Pakistani men (Table 4).

**Table 4.** Mean relative abundance of major genera (detected in 100% gut samples) and their comparison to the uBiome dataset (see Table S5 for complete list).

| Genus | Mean relative abundance (all) |  |  | Mean relative abundance (males) |  |  | Mean relative abundance (females) |  |  | uBiome<sup>a</sup> |
| --- | --- | --- | --- | --- | --- | --- | --- | --- | --- | --- |
|  | *min* | *mean* | *max* | *min* | *mean* | *max* | *min* | *mean* | *max* |  |
| *Faecalibacterium* | 1.62 | 8.89 | 21.20 | 2.39 | 7.31 | 12.77 | 1.62 | 10.19 | 21.20 | 13.11 |
| *Bacteroides* | 0.28 | 7.91 | 42.25 | 0.28 | 3.76 | 13.50 | 0.28 | 11.33 | 42.25 | 22.71 |
| *Blautia* | 0.48 | 4.85 | 15.55 | 1.49 | 3.86 | 10.86 | 0.48 | 5.66 | 15.55 | 11.43 |
| *Bifidobacterium* | 0.03 | 4.21 | 11.27 | 0.03 | 3.69 | 10.62 | 0.38 | 4.65 | 11.27 | 2.08 |
| *Pseudobutyrvibrio* | 0.38 | 3.82 | 10.41 | 0.38 | 3.29 | 10.12 | 0.52 | 4.26 | 10.41 | 2.64 |
| *Sarcina* | 0.25 | 3.64 | 11.49 | 0.25 | 2.82 | 6.58 | 0.27 | 4.31 | 11.49 | 2.10 |
| *Roseburia* | 0.48 | 3.46 | 8.54 | 0.48 | 2.85 | 5.03 | 1.05 | 3.96 | 8.54 | 7.75 |
| *Sutterella* | 0.13 | 3.15 | 9.98 | 0.13 | 2.49 | 7.41 | 0.21 | 3.69 | 9.98 | 1.21 |
| *Collinsella* | 0.25 | 2.06 | 6.39 | 0.25 | 1.61 | 6.06 | 0.63 | 2.43 | 6.39 | 1.88 |
| *Subdoligranulum* | 0.28 | 2.00 | 4.10 | 0.28 | 1.53 | 3.77 | 0.34 | 2.38 | 4.10 | 2.93 |
| *Dorea* | 0.35 | 1.95 | 5.06 | 0.35 | 1.61 | 5.06 | 0.38 | 2.23 | 5.01 | 1.46 |
| *Alistipes* | 0.06 | 0.92 | 6.81 | 0.06 | 0.49 | 1.91 | 0.08 | 1.28 | 6.81 | 1.88 |
| *Clostridium* | 0.12 | 0.81 | 3.15 | 0.12 | 0.42 | 1.00 | 0.13 | 1.14 | 3.15 | 0.82 |
| *Fusicatenibacter* | 0.03 | 0.77 | 2.41 | 0.10 | 0.64 | 2.32 | 0.03 | 0.88 | 2.41 | 2.65 |
| *Intestinibacter* | 0.13 | 0.68 | 3.34 | 0.13 | 0.40 | 0.90 | 0.17 | 0.92 | 3.34 | 0.86 |
| *Lachnospira* | 0.06 | 0.41 | 1.37 | 0.06 | 0.22 | 0.55 | 0.10 | 0.56 | 1.37 | N/A |
| *Intestinimonas* | 0.01 | 0.20 | 0.81 | 0.01 | 0.14 | 0.25 | 0.04 | 0.25 | 0.81 | N/A |

<sup>a</sup>Numbers as reported on uBiome Explorer in July 2018. Numbers are subject to change with inclusion of more healthy participants to the ongoing uBiome citizen science initiative (e.g. [41]).

*Faecalibacterium* are amongst the most commonly detected bacteria in the human gut that break down complex and resistant starches (e.g. legumes and whole grains). There is mixed evidence regarding their roles in protecting humans from inflammatory bowel diseases [62,63]. Compared to uBiome dataset, *Faecalibacterium* were under-represented in Pakistani males (7.3% vs. 13.11%) (Table 4) and its abundance was positively correlated with *Subdoligranulum* (τ = 0.45, Figure S4, Table S6). *Bacteroides* are more commonly observed in people on protein and animal fat diet [61]. *Bacteroides* were significantly under-represented in Pakistani males and females comprising only 3.75% and 11.33% of the total bacterial population respectively compared to roughly 23% in the healthy individuals (Table 4). Bacteroides were negatively correlated with *Alloprevotella* (τ = −0.48, Figure S4, Table S6). *Roseburia* is another important component of the healthy gut microbiota. These organisms produce butyrate that has anti-inflammatory activities and might even prevent colon cancer [64]. *Roseburia* comprised of, on average, 2.85% and 3.95% of total bacterial population in Pakistani males and females, respectively, which was lower than 7.75% typically seen in uBiome healthy individuals (Table 4). *Blautia* also help digest complex carbohydrates and their levels are increased in healthy individuals compared to people with diabetes [65]. *Balutia* were also under-represented in the Pakistani individuals comprising only 3.85% and 5.66% of total genera in Pakistani males and females relative to ∼11% in the gut of healthy individuals (Figure 5A and Table 3). Its abundance was positively correlated with several beneficial bacteria such as *Dorea* (τ = 0.65), *Marvinbryantia* (0.55), *Subdoligranulum* (0.55), *Collinsella* (0.52), *Fusicatenibacter* (0.51), *Intestinimonas* (0.5), and *Clostridium* (0.49) (Figure S4 and Table S6).

### High Levels of Probiotics in the Pakistani Gut

Despite obvious concerns of high load of Proteobacteria (*Succinivibrio*) and a skewness towards Firmicutes (fat digesting microorganisms) in the Pakistani gut, we detected high prevalence and abundance of beneficial microorganisms such as *Bifidobacterium* (31/31 samples) and *Lactobacillus* (30/31) in the Pakistani gut (Table S5). *Bifidobacterium* are amongst the most well-studied microorganisms in the human gut [66]. They are especially abundant in infants where they help digest indigestible oligosaccharides and fiber from breast milk and may protect gut from inflammation [66]. Due to their usefulness, many *Bifidobacterium* species are sold as over-the-counter probiotics. *Bifidobacterium* was the fourth most abundant genus in the Pakistani gut comprising 4.21% of the total community (males = 3.68% and females = 4.65%) relative to 2.08% in uBiome dataset (Table 4). Similarly, *Lactobacillus* is also commercially sold as probiotics and was detected in 31/32 Pakistani samples, albeit with mean relative abundance of <1% (Table S5). The relatively high levels of probiotics in the Pakistani gut could be due to the fact that milk-based products and fermented foods (e.g. yoghurt) are very popular in the Pakistani diet, especially in Punjab. Moreover, infants are typically breastfed for up to two years for religious and cultural reasons throughout Pakistan (20/29 of our survey respondents, Table S1). Previously, an association between *Bifidobacterium* and milk-based diets has been established [67]. The most notable absentee from the majority of Pakistani samples, however, was *Akkermansia* (Verrumicrobia) that has many health benefits in the human body [68]. *Akkermansia* has been associated with a reduced risk of obesity, type 2 diabetes, and also combats weight gain and inflammation [68]. However, it was one of the low-count and low-variance features removed during our analysis (see Materials and Methods). In the original pre-filtered dataset, *Akkermansia* was detected in 2/32 samples at very negligible amounts (<0.06%) and in one sample (ISB-18) at 2.76% (data not shown). Finally, in addition to gender, *Mogibacterium* genus was differentially abundant in people with different social habits (Figure S5), although we did not explore this further.

### The Composition of the Oral Microbiome

Pakistani oral microbiome (*n* = 28 samples) was dominated by five phyla constituting ∼99% of the total microbial community (Figure 6A and Table 5). These included Proteobacteria, Firmicutes, Bacteroidetes, Actinobacteria, and Fusobacteria. *Cand*. Saccharibacteria (formerly TM7), Spirochaetes, and Synergistes were also detected, albeit in minor amounts (<2% cumulative) (Table S8). Once again, gender appeared to be the strongest determinant of microbiome variability among Pakistani individuals. Similar to the gut microbiota profile, Firmicutes were significantly more abundant in women (36.28% vs. 25.43%, Figures 6B and 6C). Compared to the uBiome dataset, Pakistani individuals were over-represented in all bacterial phyla except Firmicutes (31.24% vs. 57.36%, Figure 6B and Table 5). Following quality control and filtering (see Materials and Methods), 58 genera were detected in the oral samples (Table S9). Among these, 18 were detected in all samples and included notable inhabitants of the oral cavity such as *Streptococcus, Neisseria, Haemophilus, Veillonella, Prevotella*, and *Gemella* (Table 6). While the mean relative abundances of these genera varied considerably among males and females, five genera (*Streptococcus* [Firmicutes], *Haemophilus* [Proteobacteria], *Granulicatella* [Firmicutes], *Eikenella* [Proteobacteria], and *Leptotrichia* [Fusobacteria]) had significant differential abundance in females and males (Figure 6C and Table S10). Of these, *Streptococcus, Haemophilus*, and *Granulicatella* were significantly more abundant in females versus males (*FDR* < 0.05) while *Eikenella* and *Leptotrichia* were significantly more abundant in males versus females (Figure 6C).

**Table 5.** Mean relative abundance of major phyla (detected in 100% oral samples) and their comparison to the uBiome dataset (see Table S8 for complete list).

| **Phylum** | **Mean relative abundance (all)** |  |  | **Mean relative abundance (males)** |  |  | **Mean relative abundance (females)** |  |  | **uBiome<sup>a</sup>** |
| --- | --- | --- | --- | --- | --- | --- | --- | --- | --- | --- |
|  | *min* | *mean* | *max* | *min* | *mean* | *max* | *min* | *mean* | *max* |  |
| Proteobacteria | 8.28 | 34.66 | 74.85 | 8.28 | 33.89 | 51.32 | 24.88 | 35.33 | 74.85 | 23.18 |
| Firmicutes | 15.03 | 31.25 | 62.43 | 15.03 | 25.43 | 42.99 | 16.10 | 36.29 | 62.43 | 57.36 |
| Bacteroidetes | 4.13 | 15.43 | 31.10 | 5.12 | 16.95 | 31.10 | 4.13 | 14.12 | 21.61 | 10.93 |
| Actinobacteria | 2.07 | 10.23 | 37.96 | 4.71 | 13.37 | 37.96 | 2.07 | 7.51 | 15.26 | 5.57 |
| Fusobacteria | 0.64 | 7.27 | 16.77 | 2.29 | 9.42 | 16.77 | 0.64 | 5.40 | 10.68 | 3.19 |

<sup>a</sup>Numbers as reported on uBiome Explorer in July 2018. Numbers are subject to change with inclusion of more healthy participants to the ongoing uBiome citizen science initiative (e.g. [41]).

**Table 6.** Mean relative abundance of major genera (detected in 100% oral samples) and their comparison to uBiome dataset (see Table S9 for complete list).

| **Genus** | **Mean relative abundance (all)** |  |  | **Mean relative abundance (males)** |  |  | **Mean relative abundance (females)** |  |  | **uBiome<sup>a</sup>** |
| --- | --- | --- | --- | --- | --- | --- | --- | --- | --- | --- |
|  | *min* | *mean* | *max* | *min* | *mean* | *max* | *min* | *mean* | *max* |  |
| *Haemophilus* | 1.00 | 16.17 | 56.21 | 1.00 | 8.25 | 16.83 | 8.33 | 23.04 | 56.21 | 13.01 |
| *Streptococcus* | 6.74 | 15.58 | 37.66 | 6.74 | 10.18 | 17.28 | 9.68 | 20.26 | 37.66 | 35.18 |
| *Neisseria* | 0.01 | 11.09 | 35.67 | 0.12 | 16.51 | 35.67 | 0.01 | 6.39 | 21.31 | 7.12 |
| *Veillonella* | 1.18 | 7.29 | 21.22 | 2.21 | 7.48 | 18.26 | 1.18 | 7.13 | 21.22 | 9.23 |
| *Leptotrichia* | 0.72 | 5.63 | 16.54 | 2.48 | 7.62 | 16.54 | 0.72 | 3.90 | 10.80 | 1.83 |
| *Capnocytophaga* | 0.46 | 4.42 | 11.41 | 0.46 | 4.82 | 11.41 | 1.00 | 4.06 | 8.90 | 0.99 |
| *Prevotella* | 0.48 | 4.16 | 23.74 | 1.03 | 5.62 | 23.74 | 0.48 | 2.89 | 8.55 | 5.09 |
| *Actinomyces* | 0.43 | 4.03 | 14.27 | 1.02 | 5.40 | 14.27 | 0.43 | 2.84 | 4.79 | 2.06 |
| *Porphyromonas* | 0.02 | 3.59 | 10.64 | 0.16 | 3.26 | 7.71 | 0.02 | 3.88 | 10.64 | 1.73 |
| *Gemella* | 0.28 | 3.42 | 18.18 | 0.73 | 1.76 | 4.24 | 0.28 | 4.86 | 18.18 | 6.91 |
| *Corynebacterium* | 0.16 | 2.77 | 11.55 | 0.16 | 3.34 | 11.55 | 0.16 | 2.28 | 5.26 | 1.34 |
| *Lautropia* | 0.03 | 2.65 | 14.28 | 0.03 | 3.40 | 10.47 | 0.03 | 2.01 | 14.28 | 1.21 |
| *Aggregatibacter* | 0.04 | 2.57 | 10.70 | 0.08 | 2.50 | 7.29 | 0.04 | 2.62 | 10.70 | 0.94 |
| *Centipeda* | 0.03 | 1.42 | 5.72 | 0.09 | 2.24 | 5.72 | 0.03 | 0.72 | 2.54 | 0.57 |
| *Granulicatella* | 0.02 | 1.03 | 3.86 | 0.02 | 0.46 | 1.97 | 0.16 | 1.51 | 3.86 | 2.00 |
| *Trichococcus* | 0.04 | 0.92 | 3.21 | 0.04 | 0.55 | 1.18 | 0.24 | 1.24 | 3.21 | N/A |
| *Bergeyella* | 0.07 | 0.86 | 3.70 | 0.07 | 0.60 | 2.17 | 0.12 | 1.09 | 3.70 | 0.52 |
| *Abiotrophia* | 0.01 | 0.64 | 3.09 | 0.01 | 0.55 | 1.57 | 0.04 | 0.73 | 3.09 | 0.78 |

<sup>a</sup>Numbers as reported on uBiome Explorer in July 2018. Numbers are subject to change with inclusion of more healthy participants to the ongoing uBiome citizen science initiative (e.g. [41]).

**Figure 6.**
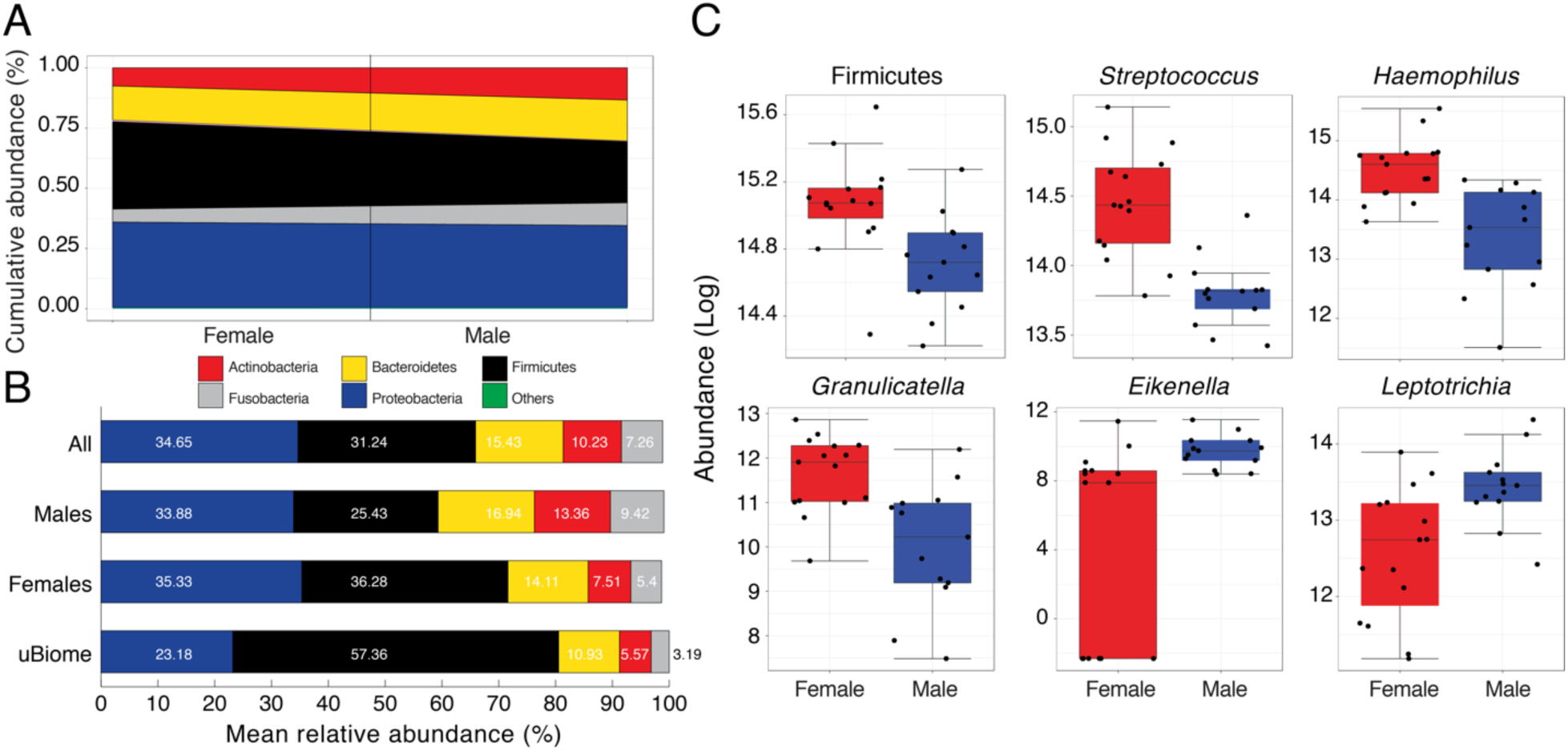
**(A)** Cumulative abundance (%) of major phyla detected in the Pakistani oral cavity. Black line separates male and female samples. *Others* category includes Spirochaetes, *Cand*. Saccharibacteria, and Synergistetes (Table S8 for actual numbers for all detected phyla) **(B)** Comparison of mean relative abundance (%) of major phyla detected in the oral cavity of Pakistani males and females versus uBiome dataset. Numbers on bars indicate actual percentages. **(C)** Boxplots comparing phyla and genera with significant differential abundance in males and females (Wilcoxon rank-sum test, *FDR* < 0.05)

*Haemophilus* and *Streptococcus* are commensal to the human body [69] although some strains can be opportunistic pathogens [70]. Both were moderately correlated with *Gemella* (τ > 0.43, Figure S6, Table S11), which along with *Neisseria* and *Veillonella* are considered cornerstones of good oral hygiene. Some *Neisseria* subtypes have even been explored as possible probiotics to stave off tooth decay. Similar to *Bifidobacterium* and *Lactobacillus* in the human gut, *Neisseria* constituted a very good load in the Pakistani oral microbial community (mean relative abundance = 11.09% vs 7.12% in the uBiome dataset, Table 6). *Neisseria* were moderately correlated with *Eikenlla* (τ = 0.44). Other than gender, however, none of the other metadata were associated with differential abundance of taxa including brush frequency, dentist visit, and other basic questions related to oral hygiene (Table S1 for metadata) except antibiotic intake as child where *Slackia* was significantly over-represented in individuals who responded ‘No’ (data not shown). Overall, oral microbiome in Pakistani individuals seemed roughly more similar to the typical oral microbiome seen in other populations (Table 6). This could be because oral microbiome is relatively more resistant to long-term effects of antibiotics and can restore quickly after antibiotic administration [71].

### Widespread Dysbiosis in the Pakistani Gut Microbiota?

Finally, we combined the gut and oral datasets and studied within and between body site species-level diversity (*n* = 52 samples, gut = 27, oral = 25) using traditional diversity measures (Figure 7). Alpha-diversity measures considering both richness (observed and Chao1) and evenness (Shannon’s index) indicated that oral samples were more diverse relative to gut samples (*P* < 0.05, Figure 7A). This was true regardless of whether we rarefied data to the minimum library size (9,437 species/sample) or not (min = 5,848, max = 195,640, mean = 52,950 species/sample) and even when we relaxed the feature (species) inclusion criteria (from having at least 4 counts in at least 20% samples to 2 counts in 10% samples, see Materials and Methods). Correlations between alpha-diversity indicators remained strong (*r* > 0.87) for all simulations (Figure S7). Thus, low diversity of Pakistani gut microbiota is likely not due to technical reasons (i.e. subsampling, data filtering) but is perhaps indicative of widespread microbiota dysbiosis in the Pakistani gut (unhealthy eating habits, frequent antibiotic usage). However, observed, Chao1, and Shannon’s diversity indicators do not consider phylogenetic relatedness of species when evaluating sample diversity. Recent simulations have indicated that abundance-weighted diversity measures incorporating phylogenetic information can outperform traditional measures [44]. Phylogenetic entropy, which is the phylogenetic generalization of Shannon’s index [45], evaluating total diversity (i.e. at all taxonomic levels) indeed confirmed that gut and oral samples were at least equally diverse and some gut samples in fact had more diversity relative to oral samples (*P* = 0.92, Figure 7B). Furthermore, a two-dimensional (2D) principal coordinate analysis (PCoA) confirmed high dissimilarity between the two body sites (*R* = 0.84, *P* < 0.001, ANOSIM) and clustering by body-site rather than by individual (Figure 7C). Oral samples, in particular, clustered more tightly together compared to gut samples that were more widely dispersed (no apparent gender-based clustering). This again shows that oral microbiome is relatively more stable to the long-term effects of antibiotics and diet and there is greater heterogeneity from individual to individual in the gut samples [71]. Differential abundance analysis followed by LDA highlighted 25 species and their relevance in characterizing gut and oral samples (Figure 7D). Several well-known gut inhabitants (e.g. *Bifidobacterium longum, Bacteroides vulgatus, Blautia faecis*, and others) characterized gut samples whereas *Streptococcus sp 11aTha1, Neisseria flava, Gemella morbillorum*, and *Haemophilus influenzae* (among others) characterized oral samples (Table S12 for complete list).

**Figure 7.**
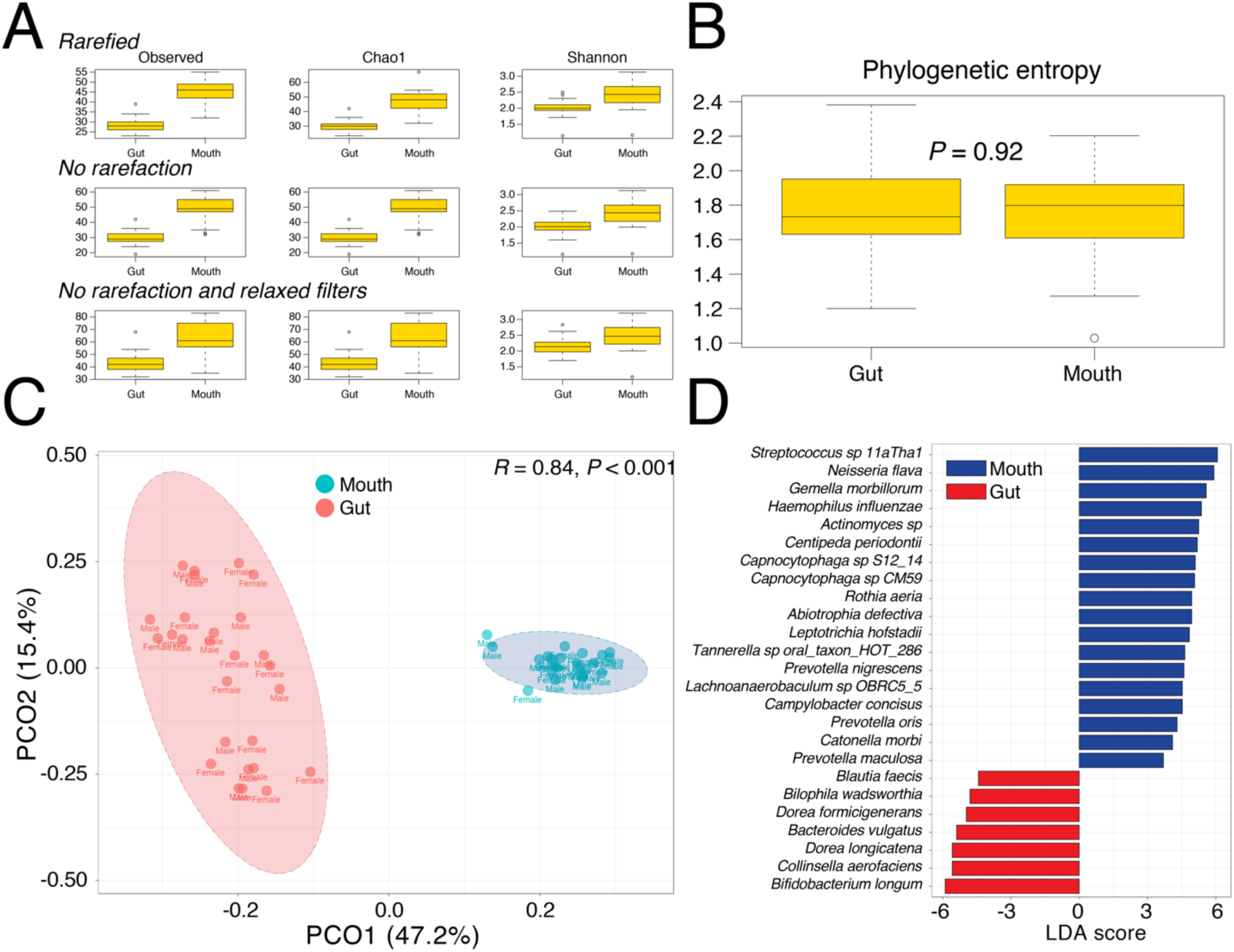
**(A)** Boxplots displaying within-sample alpha-diversity for 52 gut and oral samples (>10,000 species count/sample). These calculations were repeated after rarefying to the minimum library size, without rarefying, and relaxing the feature (species) inclusion criterion (see text). All comparisons were statistically significant (*P* < 0.05, Wilcoxon rank sum test). **(B)** Boxplot displaying within-sample alpha-diversity using phylogenetic entropy, a phylogenetic generalization of the Shannon’s index [45]. **(C)** A 2D PCoA plot evaluates dissimilarity (Weighted UniFrac) between gut and oral samples (*P* < 0.001, *R* = 0.88, ANOSIM). **(C)** Bar graph showing LDA scores of top 25 significant species characterizing gut and oral samples (see Table S12 for details).

## DISCUSSION

In this study, we describe the sequencing results of the pilot phase of the Pakistan Microbiome Initiative, a joint venture between COMSATS University Islamabad and uBiome, inc., USA. The Pakistan Microbiome Initiative, over the next two years, will target the sequencing of the gastrointestinal tract (gut) and oral cavity microbiota in ∼400-600 Pakistani adults (age >18) from the major urban and rural areas of the country. In the pilot phase, we have performed 16S rRNA gene sequencing of the gut and oral microbial communities in 32 Pakistani adults (61 total samples). These individuals were ‘cherry-picked’ to cover the maximum possible geographic and ethnic diversity of Pakistan in the early phases of the project (Table S1).

We discovered that the Pakistani gut microbiota had a relatively moderate healthy classification or wellness match compared to the uBiome dataset. Wellness match, as defined by uBiome Explorer, is the similarity/overlap in the gut microbiome composition between samples of interest and carefully selected individuals who self-reported to be in excellent medical condition and gave prior consent to participate and share their data for research projects (e.g. [41]). These data are available to both commercial users and uBiome grant awardees to aid in their research. It can be argued however that a lower wellness match to uBiome database may not necessarily imply bad health in Pakistani samples simply because of the stark dietary, genetic, and social differences between the Pakistani and Western populations. It also remains challenging to describe what is ‘healthy’ from not healthy [72]. Therefore, our statements about the healthy state of Pakistani microbiota are not definitive. A closer dissection of the Pakistani dietary and social habits however does indicate that Pakistanis, especially men, do not typically eat or live very healthy, which might explain some of the gender-based differences we observed.

In general, Pakistani women more closely resembled the gut microbiome profile of uBiome dataset relative to Pakistani men and hence, tentatively, could be referred to as ‘healthier’ relative to Pakistani men, according to the definitions above. While the Pakistani gut microbiome was largely dominated by four phyla, Firmicutes, Bacteroidetes, Proteobacteria, and Actinobacteria, that comprised, on average, ∼96% of total bacteria in both males and females, women had significantly more Firmicutes than men and men had significantly more Proteobacteria than women (Figures 2-4). This raises two important concerns. First, Pakistani gut microbiota is skewed towards Firmicutes (similar to uBiome dataset rather than South Asian countries), which has been associated with obesity [56]. The similarity to uBiome dataset might be due to favoring a meat-based diet in Pakistan (similar to Western countries) in contrast to mostly plant-based diet (as in India). Since we restricted our sampling to only young, early-career individuals (mean age = 23.4 years) with normal BMI (18-25 Kg/m^2^), we tentatively speculate that these individuals might transit from a lean body shape to overweight/obese body shape sometime later in the life. Therefore, a diet enriched in berries and cherries is recommended to increase the population of Bacteroidetes in Pakistani adults since these bacteria are known to combat weight gain [52,53]. Second, high load of Proteobacteria in gut (>13%) has previously been associated with metabolic disorders, inflammation and cancer, and considered a diagnostic marker for microbiota dysbiosis and disease [33]. Proteobacteria, on average, comprise 4.5% of the total bacterial population in healthy human gut [33]. In the Pakistani population, however, Proteobacteria comprised, on average, ∼15% of the total bacterial population (∼25% in males). The most abundant proteobacterial genus was *Succinivibrio* (mean = ∼23% in males vs. ∼2% in females, Table S5) that has previously been associated with starch metabolism [59] and is an integral member of the honey beet gut microbiota [58]. We therefore speculate that the high load of Proteobacteria in Pakistani population is likely because of consuming too many carbohydrates without being calorie/exercise conscious. Indeed, a recent country-wide diabetes prevalence survey in Pakistan revealed that a staggering one-fourth of the Pakistani population over 20 is already or may become diabetic in the near-future [73]. These are issues of genuine concern that need to be addressed immediately at the national level. We have therefore planned a time series monitoring of gut proteobacterial levels in Pakistani men to confirm whether proteobacteria are indeed chronically enriched in the Pakistani gut or not and whether it has serious health consequences.

Despite the two obvious concerns (i.e. skewness towards Firmicutes in women versus greater than expected levels of Proteobacteria in men), Pakistani gut microbial community harbored high levels of beneficial bacteria including probiotics such as *Bifidobacterium* and *Lactobacillus*, generally associated with good health. These ‘good’ bacteria are enriched in the Pakistani gut likely because of frequent consumption of milk-based products and fermented foods such as yoghurt and yoghurt drinks throughout Pakistan.

We now highlight some challenges we encountered during the initial sampling and some limitations of the present analysis. First, recruiting normal or healthy individuals throughout Pakistani is proving to be a bigger challenge than we initially expected. Roughly, 90% of the individuals we interviewed were not in the normal BMI range. Physical inactivity (self-reported to be up to 60% in a recent survey [27]) and poverty (leading to malnourishment in many parts of the country), along with bad cooking and dietary habits, are the likely culprits of imbalance between height and weight ratio among Pakistani individuals. In urban areas, we note that the majority of Pakistanis, especially men, are unconscious about calories intake and their body weight and shape and do not exercise regularly. In contrast, Pakistani women are under social pressure to maintain body weight and shape, especially before the wedding, and, in observation, consume fewer portions of the same food relative to Pakistani males. Thus, the potentially negative effects of Pakistani diet are less likely to be manifested in young Pakistani women relative to men and could partially explain the better wellness match of Pakistani women to uBiome healthy dataset. [27]). Second, antibiotic (mis)-use is frequent throughout Pakistan, which is now number three in the low or middle-income countries in antibiotic consumption (after India and China) [30]. The majority of participants initially interviewed had consumed antibiotics in the past six months and we relaxed the criteria to three-months as an afterthought. Among the selected participants, ∼84% reported they were given antibiotics as children. This is another problem we need to remedy and increase awareness about in the country. Third, Pakistani way of (over)-cooking food dilutes the nutritional value of natural ingredients and Pakistanis are not very conscious of nutritional imbalance. Pakistani diet is a mixture of high-fats and high carbohydrates that are cooked in a very unhealthy manner (personal communication with expert in nutrition biology and genomics). It thus sometimes becomes difficult to establish the accurate nutritional value of Pakistani meals.

In terms of limitations, we have only focused on bacteria (and to some extent archaea) via 16S rRNA gene sequencing and have not yet explored viruses and fungi (the mycobiome), the other big players in the human body [74]. Thus, the picture of the Pakistani microbiota remains effectively incomplete. Similarly, choice of experimental and bioinformatics protocols (PCR amplification, sequencing platform, rarefaction or not) can influence downstream results. However, results generated by uBiome laboratories follow highly reproducible experimental protocols and we observed roughly similar proportions of microbial phyla with and without rarefying (Figure S8) and are thus confident that our snapshot of the Pakistani gut and oral microbiome is rather reliable. A major issue however is of sample size and distribution. The majority of included participants belong to North Pakistan (Figure 1A) and were sampled only once. Similarly, comparisons between control and cases groups are also tentative given the small sample size and high heterogeneity between the two groups. Time-series monitoring over a larger distribution and greater number of individuals is now the next step.

In summary, our exercise provides a static snapshot of the wide ethnic and geographic diversity of the country. We are now completing the second phase of the Pakistani Microbiome Initiative and targeting hundreds of individuals from the major urban and rural areas of the country to provide more accurate estimates and hypotheses thanks to intelligence gleaned from the present study. We therefore caution the readers to be careful in extrapolating our results to the entire Pakistani community and our recommendations should be taken as generic guidelines and not as medical or professional advice. The value of the present study however cannot be underestimated as it has yielded a roadmap to more improved studies in the future and has highlighted several interesting dietary and social patterns in the Pakistani community that will be invaluable for subsequent samplings. Taken together, our results provide an overview of the gut and oral microbiota composition in the general Pakistani population (young, early-career, urban adults) and are relevant enough to be considered as baseline for future studies in the country. We are hopeful that the Pakistan Microbiome Initiative can potentially improve the dietary and social habits in the Pakistani community and can also help advance microbiology and bioinformatics research in the country.

## ACKNOWLEDGEMENTS

Authors would like to thank members of the *Computational Biology and Bioinformatics Group* and *Functional Genomics* laboratory at the COMSATS University Islamabad for their support. Special thanks to Safee ullah Chaudhary, Khubaid-ur-Rehman, Amina Bibi, Wajid Khan, and Muhammd Rizwan Riaz who helped identify potential study participants. AN is grateful to Dr. Muhammad Jawad Khan (expert in nutrition biology and genomics) for his insights regarding the nutrition value and composition of the Pakistani diet and its impact on health.

## Supplementary Materials

Quality-controlled FASTQ files of 16S rRNA gene sequencing in gut and oral samples reported in this study have been deposited to figshare (10.6084/m9.figshare.7093565).

**Figure S1.**
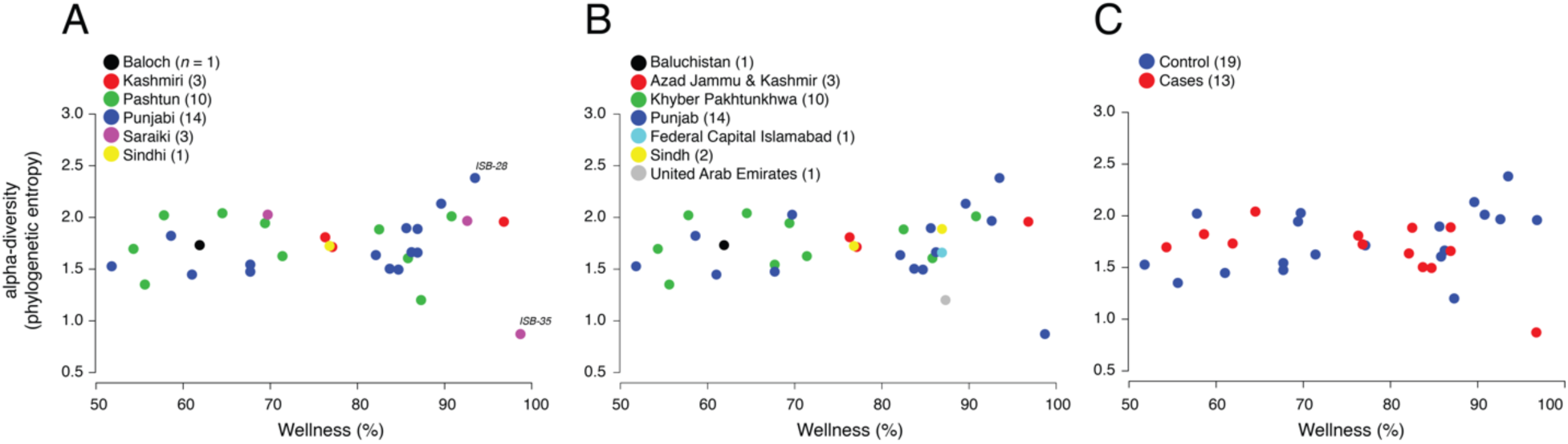
A scatter-plot displaying relationship between microbiome wellness (%) and alpha-diversity (phylogenetic entropy) for ethnicities **(A)**, geographies **(B)**, and control/cases **(C)**. AJK, Azad Jammu & Kashmir; KPK, Khyber Pakhtunkhwa.

**Figure S2.**
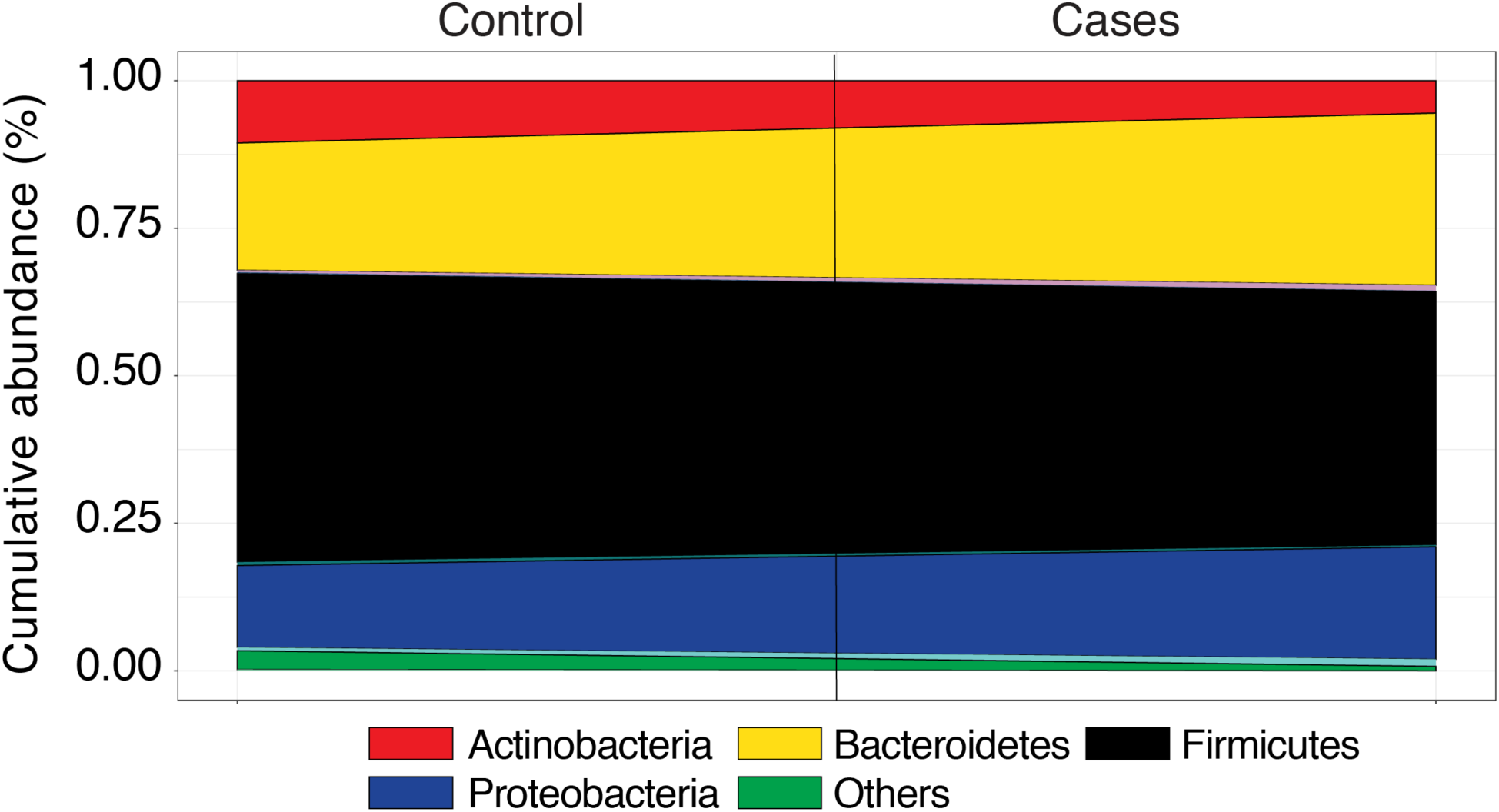
Area diagram displaying composition of microbial phyla detected in Pakistani gut samples. Black line separates control and treatment groups. *Others* category includes Tenericutes, Lentisphaerae, Euryarchaeota, Elusimicrobia, Verrucomicrobia, Fibrobacteres, Spirochaetes, and Synergistetes.

**Figure S3.**
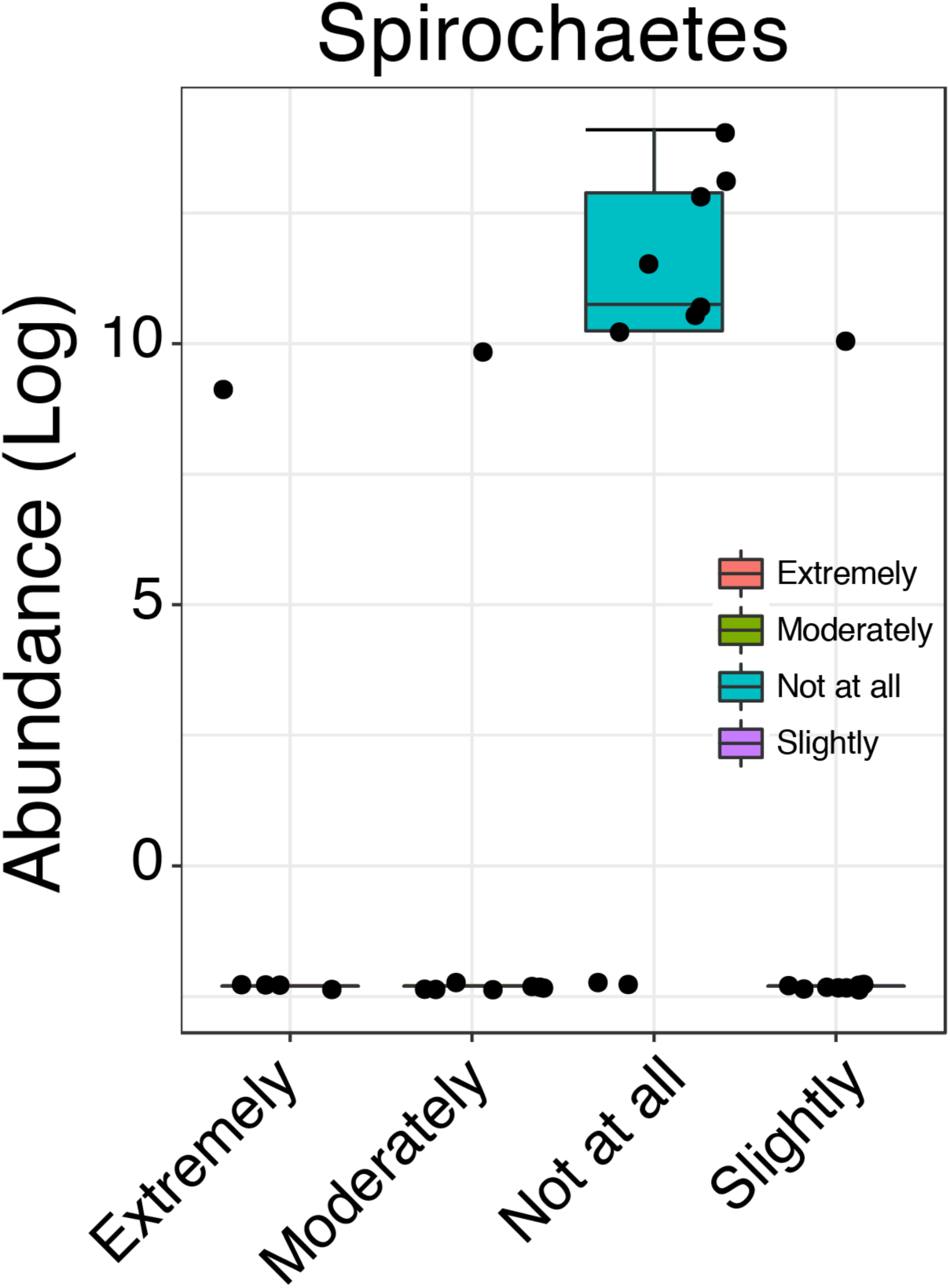
Boxplot displaying abundance distribution of Spirochaetes in four different stress states, extremely, moderately not at all, and slightly stressed. Statistical significance determined by Kruskal-Wallis test (*FDR* < 0.05).

**Figure S4.**
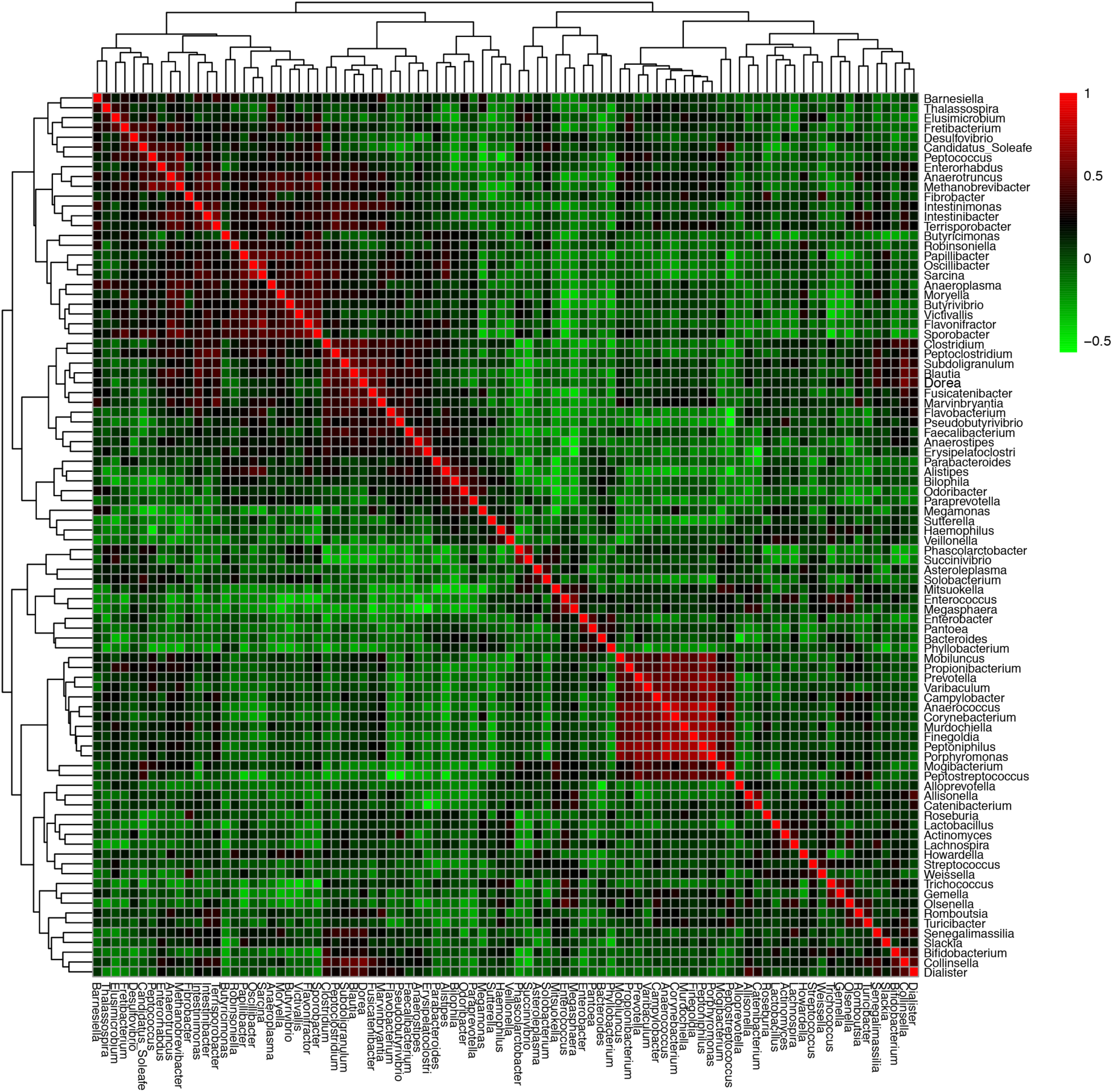
A heat-map visualizing how genera abundances in the Pakistani gut correlate to each other (Kendall’s τ). See Table S6 for actual values.

**Figure S5.**
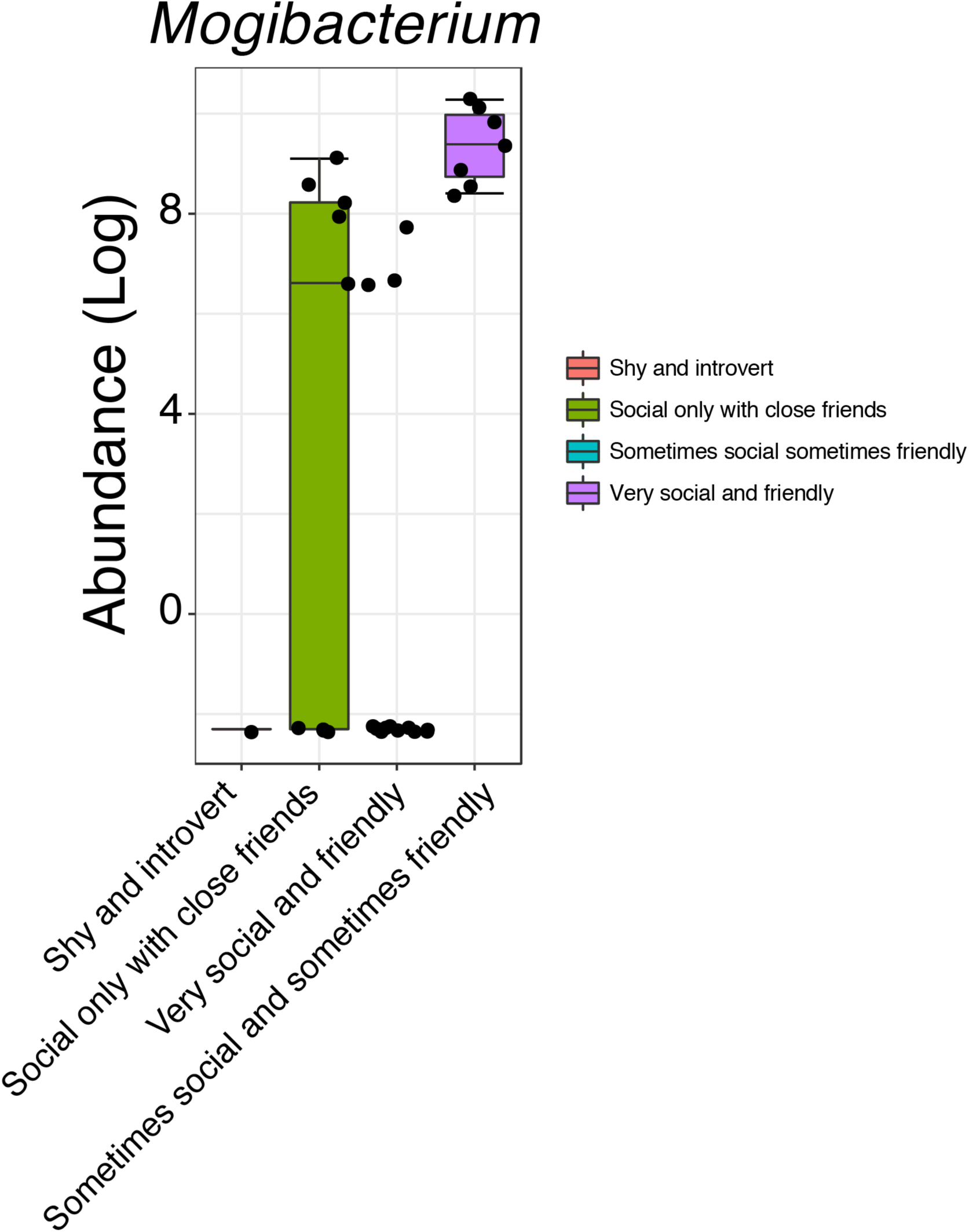
Boxplot displaying abundance distribution of *Mogibacterium* in people with different social interaction habits. Statistical significance determined by Kruskal-Wallis rank sum test (*FDR* < 0.05).

**Figure S6.**
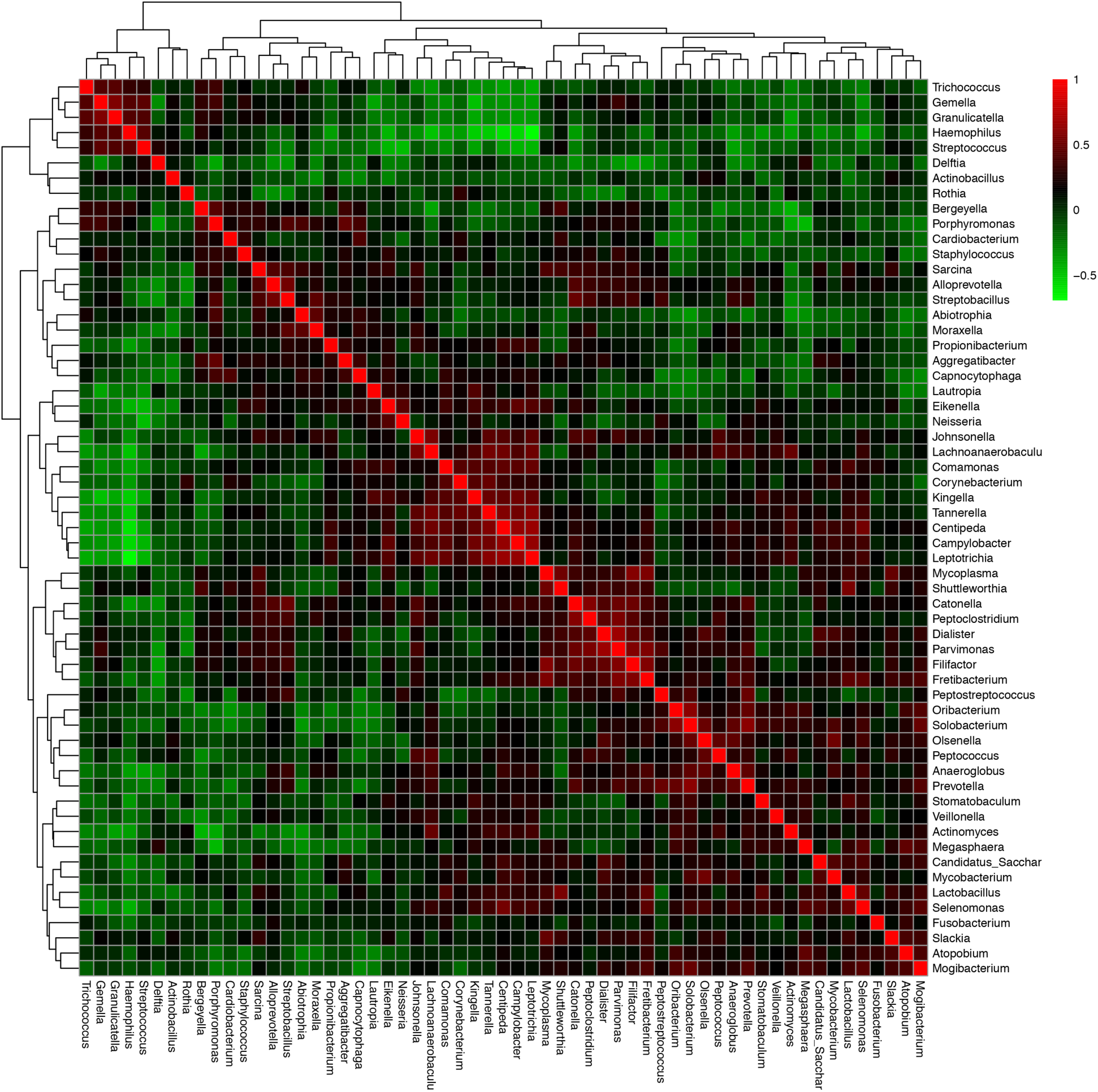
A heat-map visualizing how genera abundances in the Pakistani oral cavity correlate to each other (Kendall’s τ). See Table S11 for actual values.

**Figure S7.**
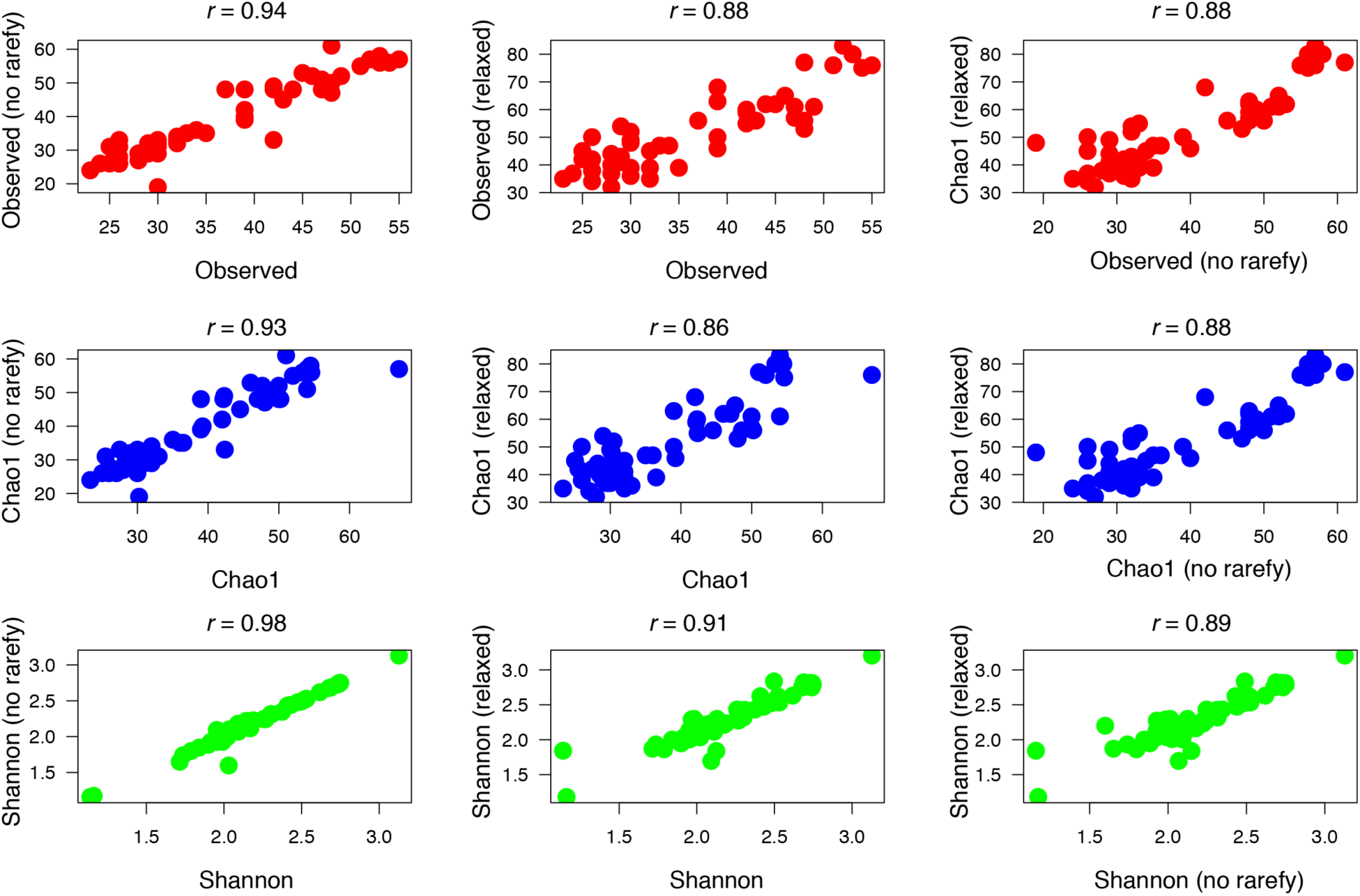
Pearson correlations (*rho*) between alpha-diversity scores for simulations described in text.

**Figure S8.**
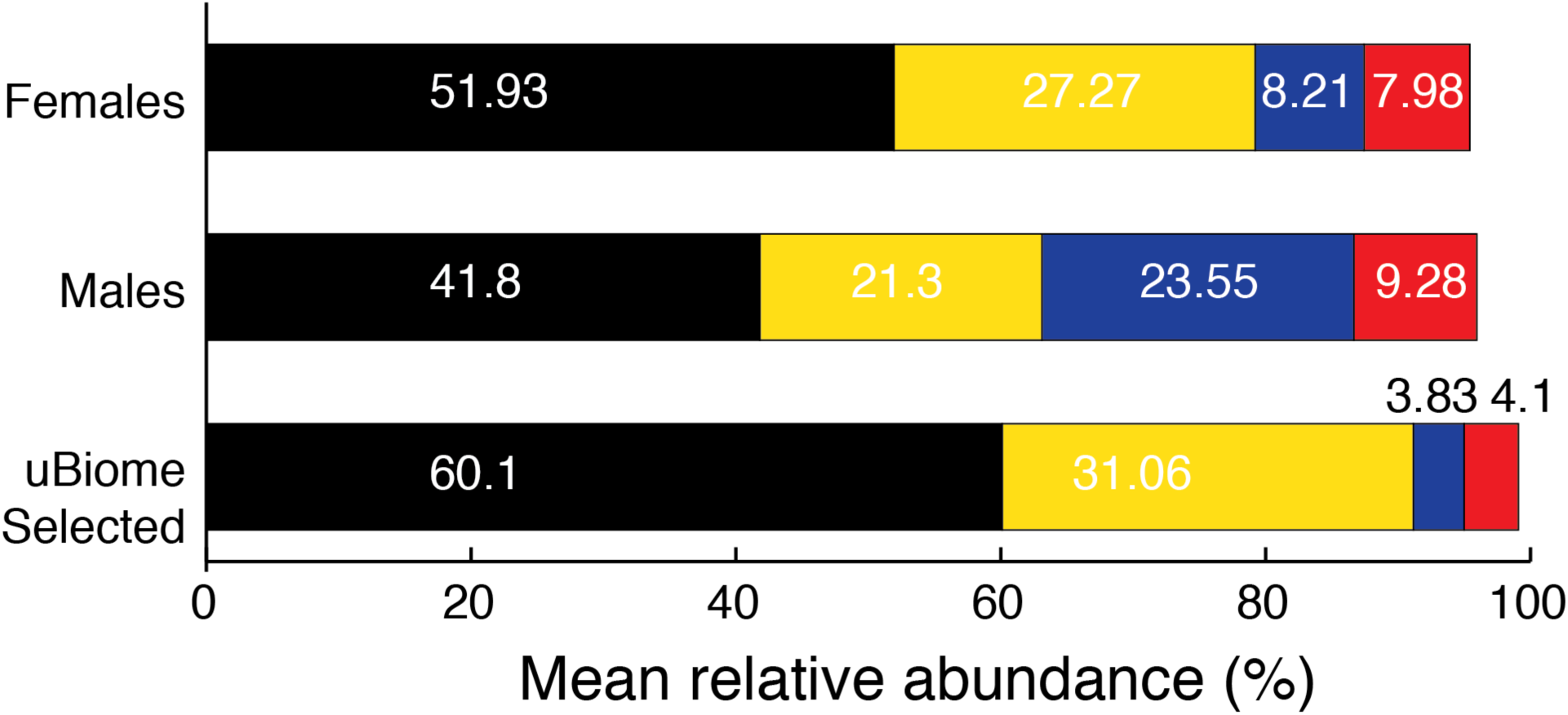
Composition of major bacterial phyla in the Pakistani gut with no subsampling (rarefying). Phyla percentages are roughly similar to Figure 2 for males and females.

**Table S1.** Detailed characteristics of shortlisted study participants and their responses to health and dietary questionnaire.

**Table S2.** Mean, minimum (*min*), and maximum (*max*) relative abundance of microbial phyla detected in all, male, and female Pakistani gut samples.

**Table S3.** Kendall’s rank (τ) abundance correlations for microbial phyla detected in Pakistani gut samples.

**Table S4.** Differential abundance analysis of microbial phyla across income groups in Pakistani gut samples. Statistically significant differences (Kruskal-Wallis test, *FDR* < 0.05) are highlighted.

**Table S5.** Mean, minimum (*min*), and maximum (*max*) relative abundance of microbial genera detected in all, male, and female Pakistani gut samples.

**Table S6.** Kendall’s rank (τ) abundance correlations for microbial genera detected in Pakistani gut samples.

**Table S7.** Differential abundance analysis of microbial genera across gender in Pakistani gut samples. Statistically significant differences (*FDR* < 0.05) are highlighted.

**Table S8.** Mean, minimum (*min*), and maximum (*max*) relative abundance of microbial phyla detected in all, male, and female Pakistani oral samples.

**Table S9.** Mean, minimum (*min*), and maximum (*max*) relative abundance of microbial genera detected in all, male, and female Pakistani oral samples.

**Table S10.** Differential abundance analysis of microbial genera across gender in Pakistani oral samples. Statistically significant differences (*FDR* < 0.05) are highlighted.

**Table S11.** Kendall’s rank (τ) abundance correlations for microbial genera detected in Pakistani oral samples.

**Table S12.** Differential abundance analysis of microbial species across body sites in Pakistani samples.

## REFERENCES

1. Conrad R, Vlassov A V. The human microbiota: composition, functions, and therapeutic potential. Med Sci Rev. 2015;2: 92–103.

2. Sender R, Fuchs S, Milo R. Are we really vastly outnumbered? revisiting the ratio of bacterial to host cells in humans. Cell. 2016;164: 337–340.

3. Shreiner AB, Kao JY, Young VB. The gut microbiome in health and in disease. Curr Opin Gastroenterol. 2015;31: 69–75.

4. Cho I, Blaser MJ. The human microbiome: at the interface of health and disease. Nat Rev Genet. 2012;13: 260–70.

5. Kaoutari A El, Armougom F, Gordon JI, Raoult D, Henrissat B. The abundance and variety of carbohydrate-active enzymes in the human gut microbiota. Nat Rev Microbiol. 2013;11: 497–504.

6. Hooper L V., Littman DR, Macpherson AJ. Interactions between the microbiota and the immune system. Science. 2012;336: 1268–1273.

7. Kau AL, Ahern PP, Griffin NW, Goodman AL, Gordon JI. Human nutrition, the gut microbiome and the immune system. Nature. 2011;474: 327–36.

8. Rath D, Amlinger L, Rath A, Lundgren M. The CRISPR-Cas immune system: Biology, mechanisms and applications. Biochimie. 2015;117: 119–128.

9. Parashar A, Udayabanu M. Gut microbiota regulates key modulators of social behavior. Eur Neuropsychopharmacol. 2016;26: 78–91.

10. Foster JA, McVey Neufeld K-A. Gut–brain axis: how the microbiome influences anxiety and depression. Trends Neurosci. 2013;36: 305–312.

11. Cryan JF, Dinan TG. Mind-altering microorganisms: the impact of the gut microbiota on brain and behaviour. Nat Rev Neurosci. 2012;13: 701–712.

12. Ley RE. Obesity and the human microbiome. Curr Opin Gastroenterol. 2010;26: 5–11.

13. Musso G, Gambino R, Cassader M. Interactions between gut microbiota and host metabolism predisposing to obesity and diabetes. Annu Rev Med. 2011;62: 361–380.

14. Evrensel A, Ceylan ME. The gut-brain axis: the missing link in depression. Clin Psychopharmacol Neurosci. 2015;13: 239–44.

15. Sommer F, Bäckhed F. The gut microbiota — masters of host development and physiology. Nat Rev Microbiol. 2013;11: 227–238.

16. Aroniadis OC, Brandt LJ. Fecal microbiota transplantation. Curr Opin Gastroenterol. 2013;29: 79–84.

17. Gilbert JA, Blaser MJ, Caporaso JG, Jansson JK, Lynch S V, Knight R. Current understanding of the human microbiome. Nat Med. 2018;24: 392–400.

18. Ursell LK, Clemente JC, Rideout JR, Gevers D, Caporaso JG, Knight R. The interpersonal and intrapersonal diversity of human-associated microbiota in key body sites. J Allergy Clin Immunol. 2012;129: 1204–1208.

19. Turnbaugh PJ, Ley RE, Hamady M, Fraser-Liggett CM, Knight R, Gordon JI. The human microbiome project. Nature. 2007;449: 804–810.

20. HMP Consortium. A framework for human microbiome research. Nature. 2012;486: 215–221.

21. Qin J, Li R, Raes J, Arumugam M, Burgdorf KS, Manichanh C, et al. A human gut microbial gene catalogue established by metagenomic sequencing. Nature. 2010;464: 59–65.

22. Bouchie A. White House unveils National Microbiome Initiative. Nat Biotechnol 2016; 34: 580.

23. Tandon D, Haque MM R. S, Shaikh S P. S, Dubey AK, et al. A snapshot of gut microbiota of an adult urban population from Western region of India. PLoS One. 2018;13: e0195643.

24. Bhute S, Pande P, Shetty SA, Shelar R, Mane S, Kumbhare S V., et al. Molecular characterization and meta-analysis of gut microbial communities illustrate enrichment of Prevotella and Megasphaera in Indian subjects. Front Microbiol. 2016;7: 660.

25. Shetty SA, Marathe NP, Shouche YS. Opportunities and challenges for gut microbiome studies in the Indian population. Microbiome. 2013; 1:24.

26. Speedy AW. Global production and consumption of animal source foods. J Nutr. 2003;133: 4048S–4053S.

27. Khuwaja AK, Kadir MM. Gender differences and clustering pattern of behavioural risk factors for chronic non-communicable diseases: community-based study from a developing country. Chronic Illn. 2010;6: 163–170.

28. Yahya F, Zafar R, Shafiq S. Trend of fast food consumption and its effect on pakistani society. Food Science and Quality Management. 2013; Available: www.iiste.org

29. Safdar NF, Bertone-Johnson E, Cordeiro L, Jafar TH, Cohen NL. Dietary patterns of Pakistani adults and their associations with sociodemographic, anthropometric and life-style factors. J Nutr Sci. 2013;2: e42.

30. Klein EY, Van Boeckel TP, Martinez EM, Pant S, Gandra S, Levin SA, et al. Global increase and geographic convergence in antibiotic consumption between 2000 and 2015. Proc Natl Acad Sci U S A. 2018;115: E3463–E3470.

31. Aziz MM, Masood I, Yousaf M, Saleem H, Ye D, Fang Y. Pattern of medication selling and self-medication practices: A study from Punjab, Pakistan. PLoS One. 2018;13: e0194240.

32. Podgorski JE, Eqani Samas, Khanam T, Ullah R, Shen H, Berg M. Extensive arsenic contamination in high-pH unconfined aquifers in the Indus Valley. Sci Adv. 2017;3: e1700935.

33. Shin N-R, Whon TW, Bae J-W. Proteobacteria: microbial signature of dysbiosis in gut microbiota. Trends Biotechnol. 2015;33: 496–503.

34. Keitel W, Petrosino J, Watson M, Dunne M. HMP Initiative 1: Human microbiome project-core microbiome sampling protocol A HMP Protocol Number: HMP-07-001. 2010. Available: http://www.fda.gov/cder/guidance/959fnl.pdf

35. Hummel W, Kula M-R. Simple method for small-scale disruption of bacteria and yeasts. J Microbiol Methods. 1989;9: 201–209.

36. Cady NC, Stelick S, Batt CA. Nucleic acid purification using microfabricated silicon structures. Biosens Bioelectron. 2003;19: 59–66.

37. Minalla AR, Dubrow R, Bousse LJ. Feasibility of high-resolution oligonucleotide separation on a microchip. In: Mastrangelo CH, Becker H, editors. 2001. pp. 90–97.

38. Mahé F, Rognes T, Quince C, de Vargas C, Dunthorn M. Swarm: robust and fast clustering method for amplicon-based studies. PeerJ. 2014;2: e593.

39. Rognes T, Flouri T, Nichols B, Quince C, Mahé F. VSEARCH: a versatile open source tool for metagenomics. PeerJ. 2016;4: e2584.

40. Glöckner FO, Yilmaz P, Quast C, Gerken J, Beccati A, Ciuprina A, et al. 25 years of serving the community with ribosomal RNA gene reference databases and tools. J Biotechnol. 2017;261: 169–176.

41. Almonacid DE, Kraal L, Ossandon FJ, Budovskaya Y V., Cardenas JP, Bik EM, et al. 16S rRNA gene sequencing and healthy reference ranges for 28 clinically relevant microbial taxa from the human gut microbiome. PLoS One. 2017;12: e0176555.

42. Weiss S, Xu ZZ, Peddada S, Amir A, Bittinger K, Gonzalez A, et al. Normalization and microbial differential abundance strategies depend upon data characteristics. Microbiome. 2017; 5:27.

43. Dhariwal A, Chong J, Habib S, King IL, Agellon LB, Xia J. MicrobiomeAnalyst: a web-based tool for comprehensive statistical, visual and meta-analysis of microbiome data. Nucleic Acids Res. 2017;45: W180–W188.

44. McCoy CO, Matsen FA. Abundance-weighted phylogenetic diversity measures distinguish microbial community states and are robust to sampling depth. PeerJ. 2013;1: e157.

45. Allen B, Kon M, Bar-Yam Y. A new phylogenetic diversity measure generalizing the shannon index and its application to phyllostomid bats. Am Nat. 2009;174: 236–243.

46. Segata N, Izard J, Waldron L, Gevers D, Miropolsky L, Garrett WS, et al. Metagenomic biomarker discovery and explanation. Genome Biol. 2011;12: R60.

47. Knights D, Ward TL, McKinlay CE, Miller H, Gonzalez A, McDonald D, et al. Rethinking “Enterotypes.” Cell Host Microbe. 2014;16: 433–437.

48. Jeffery IB, Claesson MJ, O’Toole PW, Shanahan F. Categorization of the gut microbiota: enterotypes or gradients? Nat Rev Microbiol. 2012;10: 591–592.

49. Eckburg PB, Bik EM, Bernstein CN, Purdom E, Dethlefsen L, Sargent M, et al. Diversity of the human intestinal microbial flora. Science. 2005;308: 1635–1638.

50. Harris K, Kassis A, Major G, Chou CJ. Is the Gut Microbiota a New Factor Contributing to Obesity and Its Metabolic Disorders? J Obes. 2012;2012: 1–14.

51. Duncan SH, Lobley GE, Holtrop G, Ince J, Johnstone AM, Louis P, et al. Human colonic microbiota associated with diet, obesity and weight loss. Int J Obes. 2008;32: 1720–1724.

52. Turnbaugh PJ, Hamady M, Yatsunenko T, Cantarel BL, Duncan A, Ley RE, et al. A core gut microbiome in obese and lean twins. Nature. 2009;457: 480–484.

53. Kallus SJ, Brandt LJ. The intestinal microbiota and obesity. J Clin Gastroenterol. 2012;46: 16–24.

54. De Filippo C, Cavalieri D, Di Paola M, Ramazzotti M, Poullet JB, Massart S, et al. Impact of diet in shaping gut microbiota revealed by a comparative study in children from Europe and rural Africa. Proc Natl Acad Sci U S A. 2010;107: 14691–6.

55. Conlon MA, Bird AR. The impact of diet and lifestyle on gut microbiota and human health. Nutrients. 2014;7: 17–44.

56. Ley RE, Bäckhed F, Turnbaugh P, Lozupone CA, Knight RD, Gordon JI. Obesity alters gut microbial ecology. Proc Natl Acad Sci U S A. 2005;102: 11070–5.

57. Petri RM, Schwaiger T, Penner GB, Beauchemin KA, Forster RJ, McKinnon JJ, et al. Changes in the rumen epimural bacterial diversity of beef cattle as affected by diet and induced ruminal acidosis. Appl Environ Microbiol. 2013;79: 3744–3755.

58. Mattila HR, Rios D, Walker-Sperling VE, Roeselers G, Newton ILG. Characterization of the active microbiotas associated with honey bees reveals healthier and broader communities when colonies are genetically diverse. PLoS One. 2012;7: e32962.

59. O’Herrin SM, Kenealy WR. Glucose and carbon dioxide metabolism by Succinivibrio dextrinosolvens. Appl Environ Microbiol. 1993;59: 748–55.

60. Kiefer I, Rathmanner T, Kunze M. Eating and dieting differences in men and women. J Men’s Heal Gend. 2005;2: 194–201.

61. Wu GD, Chen J, Hoffmann C, Bittinger K, Chen Y-Y, Keilbaugh SA, et al. Linking long-term dietary patterns with gut microbial enterotypes. Science. 2011;334: 105–8.

62. Jia W, Whitehead RN, Griffiths L, Dawson C, Waring RH, Ramsden DB, et al. Is the abundance of Faecalibacterium prausnitzii relevant to Crohn’s disease? FEMS Microbiol Lett. 2010;310: 138–144.

63. Sokol H, Pigneur B, Watterlot L, Lakhdari O, Bermúdez-Humarán LG, Gratadoux J-J, et al. Faecalibacterium prausnitzii is an anti-inflammatory commensal bacterium identified by gut microbiota analysis of Crohn disease patients. Proc Natl Acad Sci U S A. 2008;105: 16731–6.

64. Tamanai-Shacoori Z, Smida I, Bousarghin L, Loreal O, Meuric V, Fong SB, et al. *Roseburia* spp.: a marker of health? Future Microbiol. 2017;12: 157–170.

65. Egshatyan L, Kashtanova D, Popenko A, Tkacheva O, Tyakht A, Alexeev D, et al. Gut microbiota and diet in patients with different glucose tolerance. Endocr Connect. 2016;5: 1–9.

66. Schell MA, Karmirantzou M, Snel B, Vilanova D, Berger B, Pessi G, et al. The genome sequence of Bifidobacterium longum reflects its adaptation to the human gastrointestinal tract. Proc Natl Acad Sci U S A. 2002;99: 14422–7.

67. Sela DA, Mills DA. Nursing our microbiota: molecular linkages between bifidobacteria and milk oligosaccharides. Trends Microbiol. 2010;18: 298–307.

68. Cani PD, de Vos WM. Next-Generation Beneficial Microbes: The case of Akkermansia muciniphila. Front Microbiol. 2017;8: 1765.

69. Dewhirst FE, Chen T, Izard J, Paster BJ, Tanner ACR, Yu W-H, et al. The human oral microbiome. J Bacteriol. 2010;192: 5002–17.

70. Tikhomirova A, Kidd SP. *Haemophilus influenzae* and *Streptococcus pneumoniae*: living together in a biofilm. Pathog Dis. 2013;69: 114–126.

71. Zaura E, Brandt BW, Joost Teixeira De Mattos M, Buijs MJ, Caspers MPM, Rashid M-U, et al. Same exposure but two radically different responses to antibiotics: resilience of the salivary microbiome versus long-term microbial shifts in feces. MBio. 2015; 6: e01693–15.

72. Bäckhed F, Fraser CM, Ringel Y, Sanders ME, Sartor RB, Sherman PM, et al. Defining a healthy human gut microbiome: current concepts, future directions, and clinical applications. Cell Host Microbe. 2012;12: 611–622.

73. Basit A, Fawwad A, Qureshi H, Shera AS, NDSP Members N. Prevalence of diabetes, pre-diabetes and associated risk factors: second National Diabetes Survey of Pakistan (NDSP), 2016-2017. BMJ Open. 2018;8: e020961.

74. Filyk HA, Osborne LC. The multibiome: the intestinal ecosystem’s influence on immune homeostasis, health, and disease. EBioMedicine. 2016;13: 46–54.

